# Tuning levels of low-complexity domain interactions to modulate endogenous oncogenic transcription

**DOI:** 10.1101/2021.08.16.456551

**Authors:** Shasha Chong, Thomas G.W. Graham, Claire Dugast-Darzacq, Gina M. Dailey, Xavier Darzacq, Robert Tjian

## Abstract

Gene activation by mammalian transcription factors (TFs) requires dynamic, multivalent, and selective interactions of their intrinsically disordered low-complexity domains (LCDs), but how such interactions mediate transcription remains unclear. It has been proposed that extensive LCD-LCD interactions culminating in liquid-liquid phase separation (LLPS) of TFs is the dominant mechanism underlying transactivation. Here, we investigated how tuning the amount and localization of LCD-LCD interactions *in vivo* affects transcription of endogenous human genes. Quantitative single-cell and single-molecule imaging reveals that the oncogenic TF EWS/FLI1 requires a finely tuned range of LCD-LCD interactions to efficiently activate target genes. Modest or more dramatic increases in LCD-LCD interactions toward putative LLPS repress EWS/FLI1-driven transcription in patient cells. Likewise, ectopically creating LCD-LCD interactions to sequester EWS/FLI1 into a bona fide LLPS compartment, the nucleolus, inhibits EWS/FLI1-driven transcription and oncogenic transformation. Our findings reveal fundamental principles underlying LCD-mediated transcription and suggest mislocalizing specific LCD-LCD interactions as a novel therapeutic strategy for targeting disease-causing TFs.

## Main Text

Transcription control is a front-line mechanism underlying differential gene expression that drives animal development and disease. As critical regulators of mammalian gene expression, transcription factors (TFs) contain sequence-specific DNA-binding and transactivation domains that participate in specific protein-protein interactions to direct gene transcription. DNA-binding domains usually have well-defined protein structures and recognize specific DNA sequences in a “lock-and-key” manner with precise stoichiometries *(1, 2)*. In contrast, transactivation domains typically contain low-complexity sequence domains (LCDs) that are intrinsically disordered and not amenable to conventional structure-function analysis. Recent advances in quantitative live-cell imaging revealed that dynamic, multivalent, and selective interactions occur between various TF LCDs and other intrinsically disordered regions found in the RNA Polymerase II (Pol II) transcription machinery and associated cofactors *(3–7)*. These multivalent interactions help enrich TFs and other transcription regulators at specific genomic loci to form dynamic protein assemblies that we and others referred to as LCD-mediated “hubs” *(8–11)*. Under certain conditions, LCD overexpression can induce normally small transient hubs to develop properties resembling those of droplets formed via liquid-liquid phase separation (LLPS) *(8)*. In recent years, phase-separated membraneless compartments, often called “condensates” or “droplets”, have been invoked as an essential mechanism empowering many biological processes, including transcriptional regulation. Most studies thus far deploy in vitro experiments in test tubes to observe the formation of condensates and characterize their properties, yet the in vivo biophysical characteristics of condensates and their functional impact under physiological conditions remains poorly understood *(12)*. While it is clear that dynamic multivalent LCD-LCD interactions underlying both hub formation and LLPS could play a role in transactivation *(3–7, 13–15)*, a causal link between LLPS and transactivation has remained elusive.

Studies of how the transcriptional output changes upon manipulation of LCD-LCD interactions are just emerging and have raised new questions. On the one hand, recent reports suggest that light-induced LLPS of TF LCDs increases global cellular transcription as well as transcription of a reporter gene within a transiently transfected plasmid *(16, 17)*. On the other hand, transcription of a synthetic gene array increased with local TF concentration driven by multivalent interactions of TF LCDs, but LLPS of the LCDs did not further enhance expression of the same synthetic gene array *(18)*. These intriguing but conflicting findings necessitate new approaches to further investigate the functional impact of tuning LCD-LCD interactions and inducing LLPS on transcription in a physiological context. One as yet unmet challenge is to measure how transcription of specific endogenous genes is affected by manipulation of local LCD-LCD interactions to induce the formation of LLPS-like condensates at target loci. Here, we addressed this critical issue and revealed new principles underlying LCD-interaction-mediated transactivation of natural endogenous genes with quantitative single-cell and single-molecule imaging.

### Enhanced levels of EWS LCD self-interactions repress EWS/FLI1-driven transcription

We focused on the oncogenic TF EWS/FLI1, a potent driver of the oncogenic transcription program in Ewing’s sarcoma cells. EWS/FLI1 is a fusion TF consisting of a transactivating LCD from EWSR1 (EWS LCD) and a DNA-binding domain from FLI1 that specifically recognizes the GGAA sequence. Recent studies showed that EWS/FLI1 forms local high-concentration hubs at GGAA microsatellites, highly repetitive GGAA-containing elements in the genome, via both EWS/FLI1-DNA interactions and multivalent EWS LCD self-interactions *(8, 21, 22)*. The weak and transient multivalent interactions are required for EWS/FLI1 to activate transcription of GGAA-microsatellite-associated genes and to drive oncogenic transformation *(8, 22, 23)*. While abolishing or weakening EWS LCD self-interactions by mutations is known to disrupt EWS/FLI1-driven transcription *(8, 22)*, it is unknown how increasing such interactions might impact gene transcription.

We tackled this question in patient-derived cells by expressing an exogenous LCD that specifically interacts with endogenous EWS LCD, which is locally concentrated at endogenous EWS/FLI1 hubs. This strategy should lead to an increase in local LCD accumulation at EWS/FLI1 hubs and therefore GGAA-microsatellite-associated genes via multivalent LCD-LCD interactions. Specifically, we exploited a knock-in Ewing’s sarcoma cell line A673 that we recently generated by CRISPR-mediated genome editing. The endogenously expressed EWS/FLI1 is fused to a HaloTag, which can covalently bind to a fluorescent ligand and enable imaging of EWS/FLI1 at native expression levels *(8)*. We transiently expressed EWS LCD tagged with mNeonGreen (mNG) and simultaneously imaged EWS/FLI1-Halo and mNG-EWS in live cells with Airyscan confocal super-resolution microscopy (Fig. 1A). We observed that mNG-EWS evenly distributes throughout the cell nucleus with some minor heterogeneity but no detectable formation of larger discrete and droplet-like nuclear puncta often attributed to LLPS (Fig. 1A). As expected, endogenous EWS/FLI1-Halo is distributed as numerous small local high-concentration hubs in the cell nucleus (Fig. 1A). This is consistent with our previous observation *(8)* and suggests that the presence of exogenous EWS LCD does not noticeably disrupt or otherwise alter the prototypical intranuclear distribution of EWS/FLI1. We detected enrichment of exogenous EWS LCD at many individual EWS/FLI1 hubs (Fig. 1A) and clearly demonstrated such enrichment by averaging the images of > 500 hubs (Fig. 1B). By plotting the averaged radial intensity profiles of both EWS/FLI1 and EWS LCD surrounding the center of EWS/FLI1 hubs, we confirmed that the maximum concentration of exogenous EWS LCD locates at the hub center (Fig. 1B). These results suggest an increase of total local EWS LCD concentrations and EWS LCD self-interactions at GGAA-microsatellite-associated genes where endogenous EWS/FLI1 hubs are formed.

**Fig. 1.**
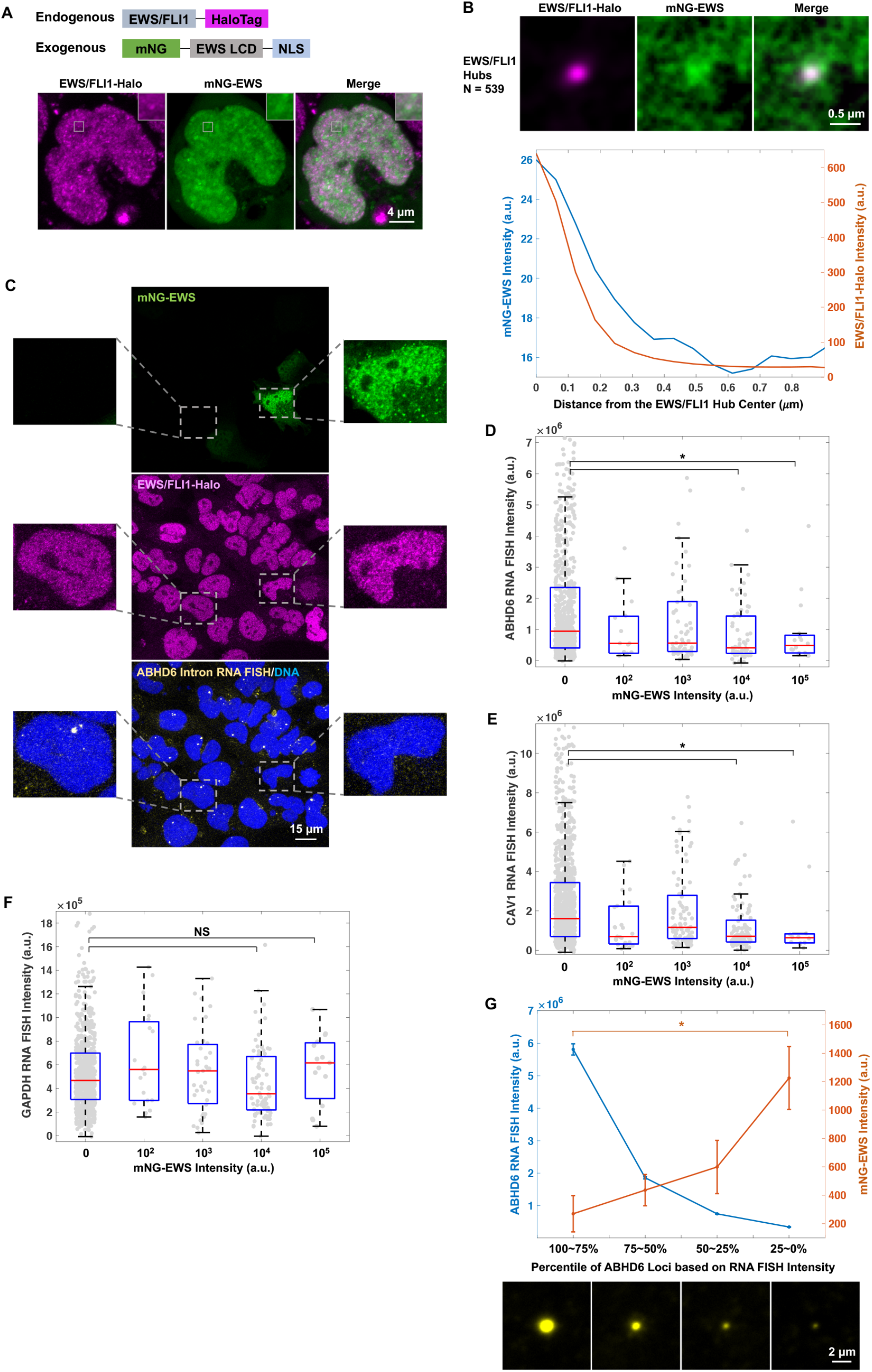
Overexpressed EWS LCD accumulates at endogenous EWS/FLI1 hubs and represses EWS/FLI1-driven transcription. (**A**) Airyscan confocal super-resolution images of endogenously expressed EWS/FLI1-Halo (JF646 labeled, magenta) and transiently expressed mNeonGreen-EWS (mNG-EWS, green) in a knock-in A673 cell. The region surrounding one particular EWS/FLI1 hub is zoomed in. mNG-EWS enrichment at the hub is visible but buried in high background. (**B**) (Upper) Averaged mNG-EWS image at 539 EWS/FLI1 hubs from 3 cells. (Lower) Radial intensity profiles of mNG-EWS (blue) and EWS/FLI1-Halo (orange) surrounding the center of EWS/FLI1 hubs on the averaged image. (**C**) Confocal fluorescence images of transiently expressed mNG-EWS (green), endogenous EWS/FLI1-Halo (JFX549 labeled, magenta), intron RNA FISH targeting *ABHD6* gene (Quasar 670 labeled, yellow), and DNA (Hoechst labeled, blue) in the knock-in A673 cells. The two cells zoomed in show that the level of *ABHD6* transcription negatively correlates with the level of mNG-EWS expression. (**D-F**) Box plots of intron RNA FISH intensities of *ABHD6* (**D**), *CAV1* (**E**), or *GAPDH* (**F**) gene after all the gene loci are categorized based on the corresponding nuclear mNG-EWS intensity. The x axis lists the order of magnitude of mNG intensities in each category. Individual gene loci are plotted in gray. *: statistically significant decrease in the RNA FISH intensity of specific mNG-positive categories compared with the mNG-negative category (p < 0.05, two-sample t-test). NS: non-significant difference between two categories. (**G**) (Upper) Averaged fluorescence intensities of intron RNA FISH (blue) and mNG-EWS (orange) at the *ABHD6* gene loci within each category. *: statistically significant increase in the mNG intensities of low-percentile *ABHD6* gene loci based on the RNA FISH intensity compared with high-percentile loci (p < 0.05, two-sample t-test). Error bars represent bootstrapped standard deviation. (Bottom) Averaged intron RNA FISH images of *ABHD6* gene loci in four categories based on the percentile rank of intron RNA FISH intensity. Each image is averaged from 211 gene loci.

To examine the effect of increasing EWS LCD self-interactions on EWS/FLI1-driven transcription, we performed simultaneous confocal imaging of transiently expressed mNG-EWS, endogenous EWS/FLI1-Halo, and intron RNA fluorescence in situ hybridization (FISH) targeting genes adjacent to GGAA microsatellites activated by EWS/FLI1, including *ABHD6* and *CAV1 (24, 25)* (Fig. 1C). Intron RNA FISH labels RNA at gene loci that are being actively transcribed, and the fluorescence intensity of a FISH punctum is proportional to the number of nascent transcripts produced from the gene locus. We took advantage of the highly variable expression levels of transiently expressed mNG-EWS to examine how transcription of GGAA-affiliated genes correlates with mNG-EWS levels within individual cells.

Specifically, we quantified the RNA FISH intensity of each gene locus and categorized all the loci based on the corresponding nuclear mNG-EWS intensity. Although within each category, RNA FISH intensities are highly variable due to the intrinsic stochasticity of gene expression *(26)*, box plots of the categories showed that transcription of *ABHD6* and *CAV1* significantly decreased with an increase in EWS LCD expression (Fig. 1D–E). As a control, no decrease of endogenous gene transcription was seen upon expression of mNG alone (Fig. S1). Moreover, transcription of *GAPDH*, a control gene not regulated by EWS/FLI1, remained unaltered regardless of EWS LCD expression (Fig. 1F). To further demonstrate the relationship between EWS LCD expression and transcription of GGAA-affiliated genes, we ranked all *ABHD6* gene loci based on their RNA FISH intensities, sorted them into four percentile-based categories, and averaged the fluorescence intensities of both RNA FISH and mNG-EWS of all the loci within each category. Across decreasing quartiles of *ABHD6* RNA FISH intensity, the average mNG-EWS intensity significantly increased (Fig. 1G), confirming that transcription of *ABHD6* decreases with increasing local EWS LCD concentration. We observed a similar inverse correlation between EWS LCD expression and transcription of another GGAA-affiliated gene *CAV1*, but not the control gene *GAPDH* (Fig. S2). Together, these results suggest that increased expression of EWS LCD leading to enhanced EWS LCD self-interactions at GGAA-affiliated genes specifically represses endogenous EWS/FLI1-driven transcription. Combining this finding with previous reports that EWS LCD self-interactions are required for EWS/FLI1 to activate transcription *(8, 22)*, we conclude that high transactivation activity requires a tightly regulated “just right” amount of EWS LCD self-interactions at endogenous target genes whereby either ablation, weakening, or increasing the interactions compromises transcriptional output. The small, transient endogenous EWS/FLI1 hubs we observed previously under native conditions achieved an optimal amount of productive LCD-LCD interactions. Here we have shown that unbalancing the system beyond this sweet spot reduces target gene expression, shedding light on the fragile stoichiometry of the system. Alternatively, however, it could be that we failed to achieve enhanced transcription because we never reached levels of LCD-LCD interactions that promoted LLPS condensate formation as has been proposed by recent reports *(16, 17)*.

### TAF15 LCD binding to EWS/FLI1 hubs induces large LLPS-like puncta and severely represses EWS/FLI1-driven transcription

We next investigated how a more potently phase-separating LCD from TAF15, which efficiently interacts with EWS LCD *(8)*, might impact the LLPS behaviors of EWS/FLI1 and EWS/FLI1-driven transcription. TAF15 LCD is known to function as a potent transactivation domain, as it multivalently interacts with intrinsically disordered regions in essential components of the transcription machinery, e.g. Pol II *(5, 8, 16)*. Moreover, TAF15 LCD was recently reported to amplify transcription in cells upon LLPS *(16)*. We simultaneously imaged transiently expressed TAF15 LCD tagged with EGFP and endogenous EWS/FLI1-Halo in the knock-in A673 cells (Fig. 2A). Whereas EGFP-TAF15 distributes throughout the cell nucleus at low expression levels, it forms discrete and very prominent droplet-like nuclear puncta at high expression levels, consistent with occurrence of what is typically classified as LLPS *(12)*. While endogenous EWS/FLI1-Halo still forms numerous small nuclear hubs in the presence of low levels of EGFP-TAF15, its nuclear distribution changes significantly upon apparent LLPS and becomes strongly enriched in the droplet-like puncta of TAF15 LCD, leaving the much smaller endogenous EWS/FLI1 hubs depleted (Fig. 2A). This result is consistent with our previous study showing heterotypic but selective interactions between TAF15 LCD and EWS LCD. We also detected such interactions at EWS/FLI1 hubs when EGFP-TAF15 is expressed at low levels by averaging the images of > 1300 hubs (Fig. 2B). We found TAF15 LCD is more enriched at the periphery instead of the center of EWS/FLI1 hubs as was observed with EWS LCD, potentially due to differences in the affinity of heterotypic TAF15-EWS LCD interactions compared with that of EWS LCD self-interactions, which we also observed in other contexts *(8)*. Nevertheless, this result confirms that at low expression levels, TAF15 LCD still binds to endogenous EWS/FLI1 hubs at GGAA-affiliated genes via heterotypic TAF15-EWS LCD interactions, increasing local LCD concentrations and therefore multivalent LCD-LCD interactions at the genes. *(8)*

**Fig. 2.**
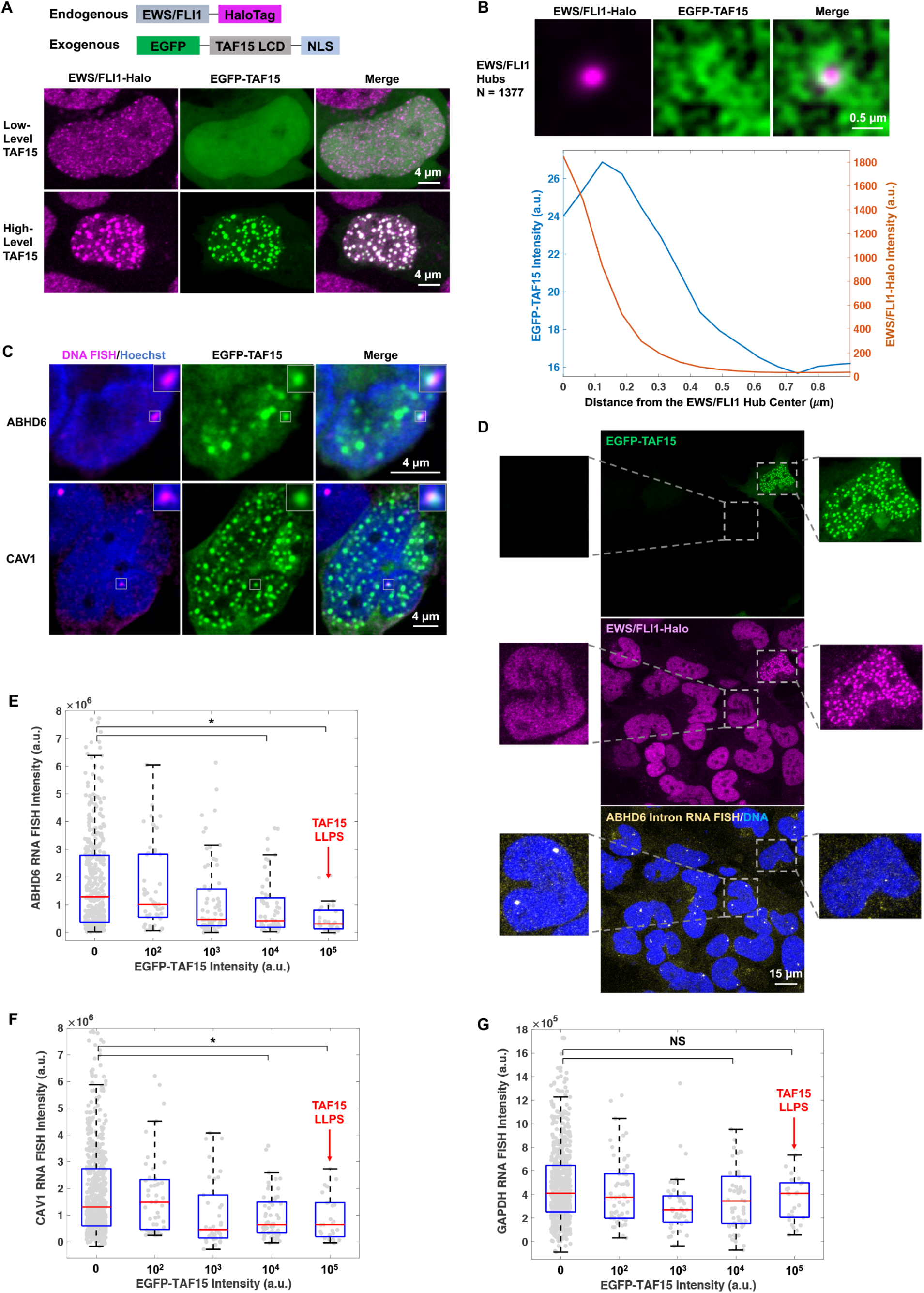
TAF15 LCD overexpression and phase separation repress EWS/FLI1-driven transcription. (**A**) Airyscan confocal super-resolution images of endogenously expressed EWS/FLI1-Halo (JF646 labeled, magenta) and transiently expressed EGFP-TAF15 (green) in two knock-in A673 cells with different levels of EGFP-TAF15. (**B**) (Upper) Averaged EGFP-TAF15 image at 1377 EWS/FLI1 hubs from 9 cells without apparent LLPS of TAF15. (Lower) Radial intensity profiles of EGFP-TAF15 (blue) and EWS/FLI1-Halo (orange) surrounding the center of EWS/FLI1 hubs on the averaged image. (**C**) Airyscan confocal super-resolution images of 3D DNA FISH targeting *ABHD6* or *CAV1* gene (enhanced Cy5 labeled, magenta), DNA (Hoechst labeled, blue), and droplet-like puncta of EGFP-TAF15 (green) in two knock-in A673 cells. The region surrounding one particular gene locus in each cell is zoomed in. Colocalization of TAF15 puncta at the loci is clearly visible. (**D**) Confocal fluorescence images of transiently expressed EGFP-TAF15 (green), endogenous EWS/FLI1-Halo (JFX549 labeled, magenta), intron RNA FISH targeting *ABHD6* gene (Quasar 670 labeled, yellow), and DNA (Hoechst labeled, blue) in the knock-in A673 cells. The two cells zoomed in show that the level of *ABHD6* transcription negatively correlates with the level of EGFP-TAF15 expression. (**E-G**) Box plots of intron RNA FISH intensities of *ABHD6* (**E**), *CAV1* (**F**), or *GAPDH* (**G**) gene after all the gene loci are categorized based on the corresponding nuclear EGFP-TAF15 intensity. The x axis lists the order of magnitude of EGFP intensities in each category. The category with apparent LLPS of TAF15 is pointed with a red arrow on each plot. Individual gene loci are plotted in gray. *: statistically significant decrease in the RNA FISH intensity of specific EGFP-positive categories compared with the EGFP-negative category (p < 0.05, two-sample t-test). NS: non-significant difference between two categories.

We next examined the spatial relationship between droplet-like TAF15 LCD puncta and GGAA-affiliated genes in cells with putative LLPS by simultaneously imaging highly expressed EGFP-TAF15 and 3D DNA FISH targeting two GGAA-affiliated genes (*ABHD6* or *CAV1*). We found 29.7% of all the detected *ABHD6* loci and 28.2% of all the detected *CAV1* loci colocalize with droplet-like TAF15 LCD puncta (Fig. 2C). *ABHD6* or *CAV1* gene-associated TAF15 LCD puncta are often slightly off-center from the gene loci, consistent with our findings in low-TAF15 cells (Fig. 2B). Our observation that not all detected GGAA target gene loci are associated with TAF15 LCD puncta naturally results from the fact that there are at most a few hundred of TAF15 LCD puncta per cell nucleus, which is orders of magnitude fewer than the ~6000 EWS/FLI1-bound GGAA microsatellites across the human genome *(28)*. Such number mismatch results in a dramatically unbalanced distribution of TAF15 LCD puncta as well as puncta-binding, endogenously expressed transcription regulators, e.g., EWS/FLI1, among all GGAA-affiliated gene loci in a cell. Whereas some of the loci are associated with puncta, having increased LCD-LCD interactions and enriching endogenous EWS/FLI1, the remaining loci are starved of TAF15 LCD as well as EWS/FLI1 that would otherwise form hubs more efficiently at the loci under native conditions.

We next examined how accumulation of TAF15 LCD at GGAA-affiliated genes and TAF15-induced LLPS affects EWS/FLI1-driven transcription with intron RNA FISH as described above for EWS LCD (Fig. 2D). Box plots showed that transcription of *ABHD6* and *CAV1* markedly decreased when TAF15 LCD expression increased. The occurrence of apparent LLPS did not reverse this trend. In fact, maximal inhibition of transactivation coincided with concentrations of exogenous TAF15 LCD required to form the prominent droplet-like puncta (Fig. 2E–F). As controls, expression of EGFP alone in these cells at comparable levels had no effect on transcription of *ABHD6* (Fig. S3) and transcription of a control gene *GAPDH* also did not track with TAF15 LCD expression (Fig. 2G), suggesting the effect of TAF15 LCD expression on endogenous transcription is specific to GGAA-affiliated EWS/FLI1 target genes. Taken together, these results suggest that allowing exogenous TAF15 LCD to accumulate at endogenous EWS/FLI1 hubs at GGAA-affiliated genes represses EWS/FLI1-driven transcription even after occurrence of TAF15-induced LLPS. Similar to increasing homotypic EWS LCD self-interactions, increasing heterotypic TAF15-EWS LCD interactions at endogenous EWS/FLI1 target genes also unbalances the sweet spot of LCD-LCD interactions needed to drive transactivation *(8, 22)*. Notably, our results are not consistent with the model where transactivation depends on the formation of LLPS condensates at target genes *(6, 7, 13, 15–17)*. Whereas the mechanism by which forced LLPS inhibits transcription of specific endogenous genes remains unclear, one possibility is that when droplet-like TAF15 LCD puncta are formed at locations other than these gene loci, the relatively stable and transcriptionally nonproductive puncta trap EWS/FLI1, thus effectively reducing the “functional pool” of EWS/FLI1 molecules that can freely access the target genes and trigger transactivation. However, because there is still a significant probability that particular GGAA-affiliated gene loci are associated with TAF15 LCD puncta (~30% for *ABHD6* and *CAV1*), it is not feasible to prove the “trapping” mechanism in this experimental setup. Besides lacking control of puncta formation locations, here we also lacked rigorous quantitative evidence for LLPS of TAF15 LCD and only determined apparent LLPS based on droplet-like behaviors *(12)*, further complicating LLPS-related conclusions. Thus, we next tested this “trapping” hypothesis and the role of LLPS in a series of experiments employing targeted mislocalization of TFs to a bona fide LLPS subcellular compartment.

### Ectopic nucleolar EWS LCD self-interactions sequester endogenous EWS/FLI1 to the nucleolus and repress EWS/FLI1-driven transcription and oncogenic transformation

Nucleoli are membraneless organelles lacking Pol II transcription and one of the best characterized examples of a condensate formed by LLPS *(19, 20)*. Here, we mislocalized LCD-LCD interactions to the nucleolus and hypothesized this might lead to the recruitment of a specific TF into the nucleolus, mimicking the above described scenario where TAF15-induced LLPS condensates formed outside the genes in question and thereby trap endogenous EWS/FLI1. This experimental setup would allow us to directly test how such trapping behavior affects transcription driven by the targeted TF. To implement such a mislocalization strategy, we fused mNG-tagged EWS LCD to a nucleolar protein, NPM1 *(29, 30)*, transiently expressed the artificial fusion protein in the knock-in A673 cells, and simultaneously imaged EWS/FLI1-Halo and mNG-EWS-NPM1 in live cells (Fig. 3A). We found mNG-EWS-NPM1 mostly localizes to the nucleolus, as expected. Interestingly, endogenous EWS/FLI1-Halo that normally localizes to the nucleoplasm is now recruited to the nucleolus. The nucleolar enrichment of EWS/FLI1-Halo calculated as the ratio of its nucleolar to nucleoplasmic concentration ([EWS/FLI1]_nucleolus_/[EWS/FLI1]_nucleoplasm_) increases with the expression level of mNG-EWS-NPM1 (Fig. S4). In contrast, EWS/FLI1-Halo remains in the nucleoplasm in cells that express a control mNG-tagged NPM1 alone (Fig. 3A, S4). These results suggest that the observed nucleolar recruitment of EWS/FLI1 by EWS-NPM1 is driven by EWS LCD self-interactions in the nucleolus. Whereas multivalent LCD-LCD interactions were thought to be weak based on the finding that they are transient *(8)* and lack the sort of well-defined structural characteristic of stoichiometric interactions, the affinity of LCD-LCD interactions was never directly compared with stoichiometric interactions in previous reports. Our results here provide a comparison between these two types of interactions *in vivo* and suggest that LCD-LCD interactions can be strong enough to outcompete stochiometric cognate TF-DNA binding interactions and trap a TF away from nucleoplasmic chromatin where it normally binds.

**Fig. 3.**
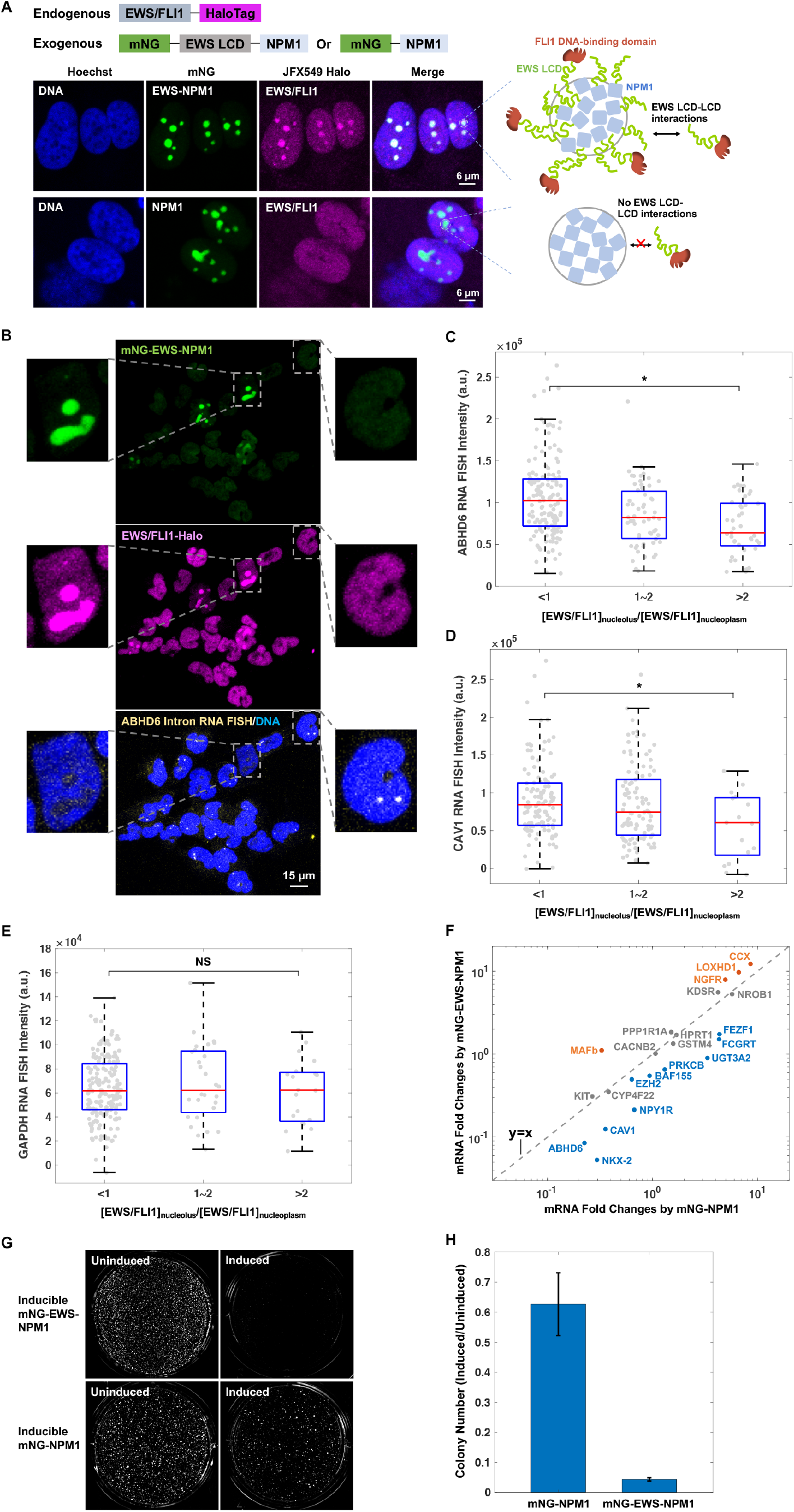
Nucleolar trapping of EWS/FLI1 represses its transcriptional activation and oncogenic transformation functions. (**A**) (Left) Airyscan confocal super-resolution images of DNA (Hoechst labeled, blue), transiently expressed mNG-EWS-NPM1 or mNG-NPM1 (green), and endogenously expressed EWS/FLI1-Halo (JF646 labeled, magenta) in the knock-in A673 cells. (Right) Schematics for protein-protein interactions within the nucleolus. (**B**) Confocal fluorescence images of transiently expressed mNG-EWS-NPM1 (green), endogenous EWS/FLI1-Halo (JFX549 labeled, magenta), intron RNA FISH targeting *ABHD6* gene (Quasar 670 labeled, yellow), and DNA (Hoechst labeled, blue) in the knock-in A673 cells. The two cells zoomed in show that the level of *ABHD6* transcription negatively correlates with the level of mNG-EWS-NPM1 expression. (**C-E**) Box plots of intron RNA FISH intensities of *ABHD6* (**C**), *CAV1* (**D**), or *GAPDH* (**E**) gene after all the gene loci are categorized based on the nucleolar enrichment of EWS/FLI1 in corresponding cells. The x axis lists the range of [EWS/FLI1]_nucleolus_/[EWS/FLI1]_nucleoplasm_ ratios for each category. Individual gene loci are plotted in gray. *: statistically significant decrease in the RNA FISH intensity of cells with nucleolar enrichment of EWS/FLI1 compared with cells without nucleolar enrichment of EWS/FLI1 (p < 0.05, two-sample t-test). NS: non-significant difference between two categories. (**F**) mRNA fold changes of 22 GGAA-affiliated EWS/FLI1 target genes by induction of mNG-EWS-NPM1 (y coordinates) versus induction of mNG-NPM1 (x coordinate) as measured by RT-qPCR. For each target gene, the mRNA level in induced cells was first normalized using the average of 4 invariant genes (Fig. S8) and then calculated as a fold change relative to the normalized mRNA level in uninduced cells. On the 2D graph, genes expressed lower upon induction of EWS-NPM1 than upon induction of NPM1 and vice versa are separated by the function plot y = x. Genes having y < x with statistical significance are plotted in blue, genes having y > x with statistical significance are plotted in orange, and genes without statistically significant difference between y and x values are plotted in gray (p < 0.05, two-sample t-test). (**G**) Soft agar colony formation assays for the knock-in A673 cells with inducible expression of mNG-EWS-NPM1 or mNG-NPM1. (**H**) Fold change of the number of colonies (diameter > 220 μm) in soft agar by induction of mNG-EWS-NPM1 or mNG-NPM1, quantified from (**G**). Error bars represent standard errors.

We next tested how nucleolar trapping of EWS/FLI1 influences its transactivation of endogenous target genes and associated oncogenic transformation functions. First, we used intron RNA FISH to measure EWS/FLI1-driven transcription at the single-cell level. Specifically, we performed simultaneous confocal imaging of transiently expressed mNG-EWS-NPM1, endogenous EWS/FLI1-Halo, and intron RNA FISH targeting GGAA-affiliated genes, *ABHD6* or *CAV1* (Fig. 3B). We quantified the RNA FISH intensity of each detected gene locus, categorized all the loci based on nucleolar mNG-EWS-NPM1 intensity or nucleolar EWS/FLI1-Halo enrichment and then generated box plots of individual categories. The box plots showed that transcription of *ABHD6* and *CAV1* significantly decreased with increasing EWS-NPM1 expression (Fig. S5, A, B, and D) and concomitant enrichment of nucleolar EWS/FLI1 (Fig. 3C–D). The decrease of endogenous gene transcription is not due to overexpression of NPM1, as expressing mNG-NPM1 did not affect transcription of *ABHD6* (Fig. S5C). Moreover, transcription of control *GAPDH* gene does not change with EWS-NPM1 expression (Fig. S5E) or nucleolar EWS/FLI1 enrichment (Fig. 3E), suggesting the effect of nucleolar trapping of EWS/FLI1 on endogenous transcription is specific to GGAA-affiliated EWS/FLI1 target genes. EWS-NPM1 could potentially be recruited to endogenous EWS/FLI1 hubs via EWS LCD self-interactions and cause repression of EWS/FLI1-driven transcription. We averaged the images of > 800 endogenous EWS/FLI1 hubs to test this “reverse recruitment” possibility, but barely detected any enrichment of EWS-NPM1 at the hubs (Fig. S6). This is likely due to the low availability of both EWS/FLI1 and EWS-NPM1 in the nucleoplasm. This finding is quite distinct from the scenario we observed above (Fig. 1B) of exogenous EWS LCD accumulating at EWS/FLI1 hubs. The lack of EWS-NPM1 accumulation at EWS/FLI1 hubs suggests that our detected repression of EWS/FLI1-driven transcription is caused by nucleolar trapping of EWS/FLI1, which effectively lowers the available concentration of EWS/FLI1 in the nucleoplasm and reduces its binding to and activation of target genes. This “repression by ectopic sequestration” represents a distinct mechanism for modulating target gene transcription from “repression via over production and LLPS induction” we observed with TAF15-induced LLPS condensates formed at target genes.

We next used a series of ensemble assays to investigate how nucleolar trapping of EWS/FLI1 affects its function in driving the oncogenic transcription program in Ewing’s sarcoma cells. To this end, we engineered the knock-in A673 cells to make a clonal cell line that expresses high levels of mNG-EWS-NPM1 upon doxycycline induction ([EWS/FLI1]_nucleolus_/[EWS/FLI1]_nucleoplasm_ ~ 1.85±0.53 (standard deviation), measured in 164 randomly chosen cells). As a control, we made a knock-in clonal cell line with inducible expression of mNG-NPM1 at levels similar to mNG-EWS-NPM1 as described above (Fig. S7). Comparing these two cell lines provides a useful way to reveal more global functional impacts of nucleolar trapping of EWS/FLI1. First, to examine how nucleolar trapping of EWS/FLI1 affects its transactivation function, we performed reverse transcription quantitative polymerase chain reaction (RT-qPCR) to measure mRNA levels of 22 GGAA-affiliated EWS/FLI1 target genes and 4 control genes not regulated by EWS/FLI1 *(8, 22)* in both cell lines before and after doxycycline induction. Whereas the control genes are expressed at constant levels regardless of the presence of exogenous EWS-NPM1 or NPM1 (Fig. S8), most GGAA-affiliated genes have transcription levels differentially affected by EWS-NPM1 versus NPM1 (Fig. 3F). Out of the 22 GGAA-affiliated genes, 45.5% are expressed significantly lower upon induction of EWS-NPM1 than upon induction of NPM1, 18.2% are expressed significantly higher upon induction of EWS-NPM1 than upon induction of NPM1, and 36.4% are equally affected by EWS-NPM1 and control NPM1. Our result suggests that many more GGAA-affiliated genes have transcription repressed rather than enhanced by nucleolar trapping of EWS/FLI1, confirming the overall repressive effect of the nucleolar trapping on EWS/FLI1-driven transcription. Next, we examined how nucleolar trapping of EWS/FLI1 affects cell proliferation by monitoring cell growth over time with the xCELLigence real-time cell analysis system *(31)*. We found that whereas the knock-in A673 cells proliferate after induction of NPM1, they completely stop proliferating 48 hours after induction of EWS-NPM1 (Fig. S9). More importantly, using soft agar colony formation assay *(32)*, we found EWS-NPM1 induction in the knock-in A673 cells abolished their malignant transformation phenotype, i.e., growth in soft agar. In stark contrast, cells after NPM1 induction continue forming colonies in soft agar, albeit with lower efficiency than uninduced cells (Fig. 3G–H). These results together suggest that nucleolar trapping of EWS/FLI1 represses the oncogenic transcription program driven by EWS/FLI1 and strongly inhibits proliferation and malignant transformation of Ewing’s sarcoma cells. Notably, the ability to tune the subcellular localization of LCD-LCD interactions thereby sequestering EWS/FLI1 and disrupting its oncogenic functions suggests a potential new therapeutic strategy for Ewing’s sarcoma, a devastating cancer still lacking effective molecular therapy.

### EWS/FLI1 diffuses more slowly in the nucleolus than in the nucleoplasm

We further investigated how nucleolar trapping of EWS/FLI1 changes its dynamic behavior. Since the nucleolus is well characterized as a LLPS condensate *(19, 20)*, measuring dynamics of EWS/FLI1 in this context provides a unique opportunity to probe how LLPS affects the dynamic behavior of a protein that localizes within condensates. We first measured the fluorescence recovery after photobleaching (FRAP) dynamics of EWS/FLI1 in the nucleolus and in the nucleoplasm, respectively. We found that FRAP dynamics of nucleolar EWS/FLI1 are significantly different from nucleoplasmic EWS/FLI1 (Fig. 4A). This suggests the two EWS/FLI1 populations differ in overall dynamics contributed by diffusion and molecular interactions *(33)*. Next, to directly quantify the diffusion dynamics of nucleolar and nucleoplasmic EWS/FLI1 in live cells, we visualized and tracked both immobile and freely diffusing molecules using stroboscopic photo-activatable single particle tracking (spaSPT) *(34–36)* (Fig. 4B). We expressed mNG-EWS-NPM1 in the knock-in A673 cells and performed spaSPT on endogenous EWS/FLI1-Halo labeled with photo-activatable Janelia Fluor 646 (PA-JF646) HaloTag ligand *(37)*. Single-molecule trajectories were classified as either nucleoplasmic or nucleolar based on binary masks generated using images of mNG-EWS-NPM1 (Fig. 4C). Nucleolar and nucleoplasmic trajectories were then separately analyzed using the Spot-On algorithm *(36)*, which fits histograms of single-particle displacements to a two-state model, where EWS/FLI1 can either be freely diffusing (‘free’), or immobile and presumably bound to protein or DNA partners (‘bound’) (Fig. 4D). Spot-On analysis yields estimates of the fraction of ‘bound’ and ‘free’ molecules as well as the apparent diffusion coefficient for each sub-population *(36)*. Interestingly, the fitted diffusion coefficient of the free subpopulation was 1.97 ± 0.20 μm^2^/s (95% bootstrap confidence interval) outside of the nucleoli but only 0.82 ± 0.26 μm^2^/s inside the nucleoli, indicating that EWS/FLI1 diffuses significantly more slowly in the nucleolus than in the nucleoplasm (Fig. 4E). This difference in free diffusion coefficient was much greater than that observed when trajectories were classified using mock nucleolar masks at randomized positions (Fig. 4E, Fig. S10), indicating that it does not result merely from geometric biases in trajectory classification. The other fit parameters from Spot-On did not differ significantly from randomized mask controls (Fig. S11). Taken together, our results are consistent with the model that the nucleolus is a true LLPS compartment displaying properties distinct from the rest of the nucleus, i.e. higher viscosity *(20, 38)*. As with endogenous nucleolar proteins *(39)*, diffusion of “guest” EWS/FLI1 molecules is reduced when they are recruited to this high-viscosity compartment by EWS-NPM1.

**Fig. 4.**
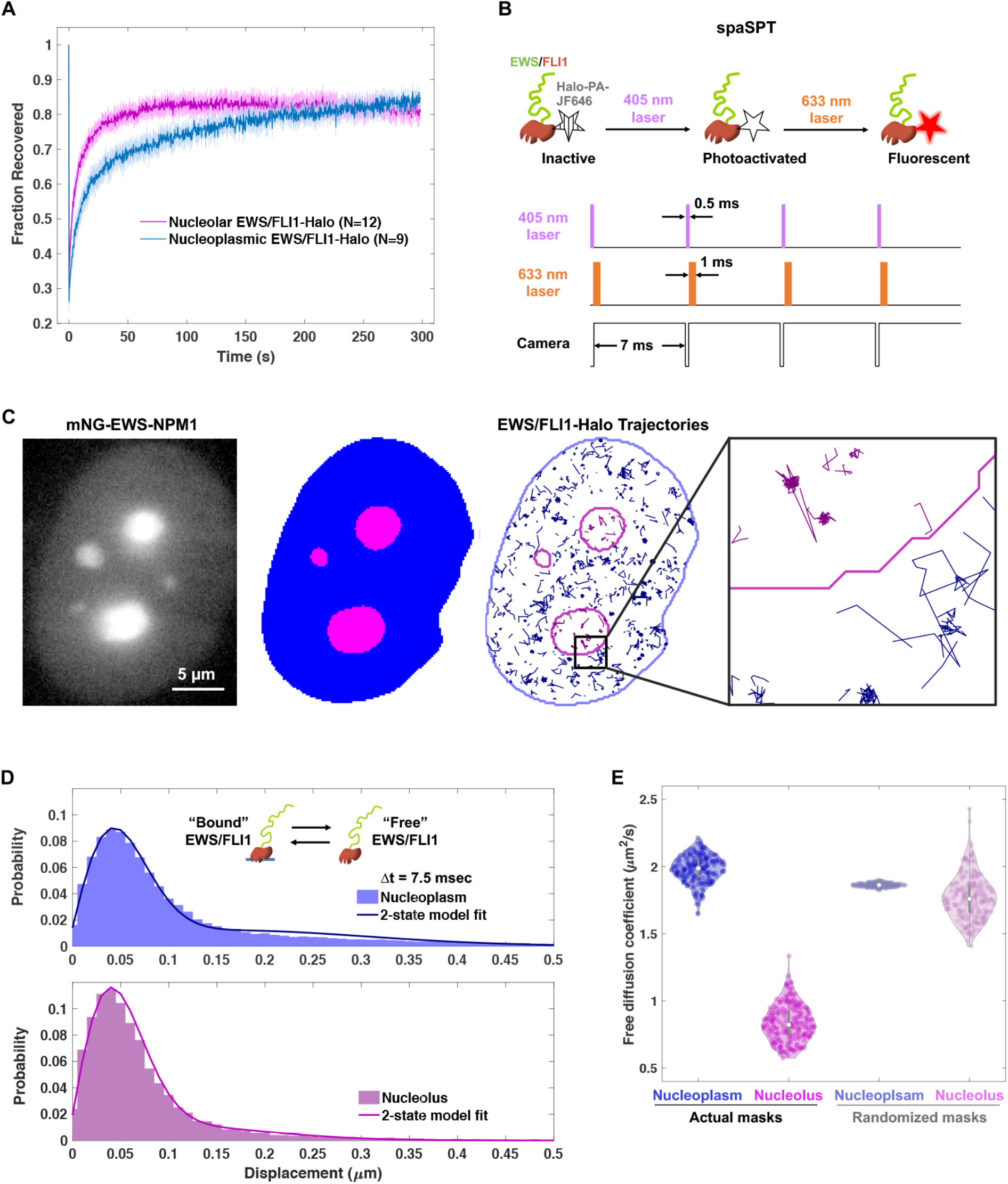
FRAP and single-molecule imaging reveal changes in the dynamic behaviors of EWS/FLI1 when it is recruited to the nucleolus. (**A**) Averaged FRAP curve of endogenous EWS/FLI1-Halo in the nucleoplasm (blue) or nucleolus (magenta). Error bars represent standard deviations. (**B**) Schematic of stroboscopic photo-activatable single particle tracking (spaSPT). Living knock-in A673 cells with endogenous EWS/FLI1-Halo expression are stained with photo-activatable HaloTag ligand PA-JF646. Short pulses of 405 nm laser and 633 nm laser are used respectively to activate and excite PA-JF646 dye to achieve ms temporal resolution and differentiate “free” molecules and “bound” molecules. (**C**) Classification of nucleoplasmic and nucleolar single-particle trajectories acquired in a spaSPT experiment. (Left) Image of mNG-EWS-NPM1 used for segmentation. Pixel intensities are displayed on a logarithmic scale between 3 and 30,000 counts. (Middle) Binary mask showing segmentation of nucleoplasm (blue) and nucleoli (magenta). (Right) Overlay of classified single-particle trajectories (thinner lines) and mask boundaries (thicker lines). Nucleoplasmic trajectories are in blue and nucleolar trajectories are in magenta. Inset: Zoom-in of a 2.6 μm square region at the boundary of a nucleolus. (**D**) Distribution histograms of displacements in a single 7.5 ms time step for trajectories in the nucleoplasm (upper, blue) and nucleolus (lower, magenta). Each histogram is fit with a two-state model. Inset shows depiction of the two-state model where EWS/FLI1 can either be freely diffusing or bound to protein or DNA partners. (**E**) Fitted diffusion coefficient of freely diffusing molecules in the nucleoplasm (blue) and nucleolus (magenta). Left 2 columns: Fits for trajectories classified using actual nucleolar masks. Violin plots show the distribution of 200 bootstrap replicates resampled by cell with replacement. Right 2 columns: Fits for trajectories classified using 200 distinct sets of randomized nucleolar masks (see Supplemental Methods).

## Discussion

Sequence-specific DNA-binding TFs have been established as key regulators of eukaryotic transcription since the early 1980s *(40)*, but how their intrinsically disordered activation domains interact with specific protein partners to direct gene activation has remained mysterious. This long-standing enigma is now starting to be unraveled with the help of recent advances in live cell single molecule imaging technologies and the discoveries of TF dynamics mediated via weak, multivalent, but selective interactions between LCDs. These emerging studies reveal that LCDs drive the formation of highly transient, small, local high-concentration TF hubs at target genomic loci that play an essential role in transactivation. Our previous data also suggest that such small transient hubs (typically containing <100 molecules with protein-binding dwell times of ~1-2 min) are quite distinct from bona fide LLPS condensates *(8)*. Excessive multivalent LCD-LCD interactions can induce the formation of LLPS-like puncta, which is usually observed under non-physiological conditions, e.g. artificial LCD over-production and/or light-induced oligomerization *(41–43)*. Originating from well-defined behaviors of polymers in solution discovered in the 19^th^ century *(44, 45)*, LLPS and condensate formation have recently reanimated excitement in biology and been proposed to underlie many cellular functions, including TF hub formation and transactivation *(46–50)*. However, studies that directly link manipulation of LCD-LCD interactions and transcriptional output are scarce *(16–18)* and the few existing studies have generated contradictory findings that stimulate important new questions. How would excessive LCD-LCD interactions and LLPS impact transactivation of endogenous genes in a physiological setting? The fact that rigorous characterization of LLPS in vivo has been challenging and often assumed or overlooked rather than experimentally confirmed *(12)* adds to the confusion of recent findings. More importantly, none of the existing studies measured transcription of specific endogenous genes upon manipulation of LCD-LCD interactions, which is essential for understanding LCD-interaction-mediated transcription in vivo under physiologically relevant contexts.

To address these knowledge gaps and to further probe mechanisms of LCD-interaction-mediated transcription, we measured transcription of endogenous target genes of the oncogenic TF EWS/FLI1 upon tuning the amount and localization of its LCD-LCD interactions in patient-derived cells. Endogenous EWS/FLI1 is known to activate genes by forming transient, small local high-concentration hubs at target gene loci via weak, multivalent EWS LCD self-interactions *(8)*. In the first part of this work, we employed quantitative single-cell imaging to study how increasing either homotypic or heterotypic LCD-LCD interactions at EWS/FLI1 target genes affects transcription output. Our findings indicate that the transactivation activity of EWS/FLI1 does not monotonically increase with the amount of local LCD-LCD interactions at genes and instead reaches a fairly narrow optimum with the endogenous amount of interactions and that further increases in LCD-LCD interactions leads to repression of transcription (Fig. 5A). This “Goldilocks principle” underlying EWS/FLI1-driven transcription – where “just the right” amount of LCD-LCD interactions is required for efficient gene activation – may be general to mammalian transcription systems and applicable to many other TFs, though the optimal amount of LCD-LCD interactions will likely vary with specific TFs and target genes. Intriguingly, we observed potent transcriptional repression of endogenous cognate genes upon apparent LLPS driven by TAF15 LCD, in contrast to previously reported increase in overall cellular transcription upon light-induced TAF15 LLPS *(16)*. We do not yet have sufficient data to provide any firm mechanistic explanation for this apparent disparity, but one possibility is related to the ability of LLPS to redistribute TFs and transcription machinery among genes via heterotypic LCD-LCD interactions (Fig. 2A, 5B). Whereas LLPS unbalances the “Goldilocks optimum” for one set of genes, it may simultaneously tune the local concentrations of transcription regulators toward the optimum for other genes. Thus, it may not be entirely unexpected to find that overall transcription is increased while specific target genes are repressed upon LLPS. Clearly, much remains to be unraveled before we can fully understand the competing functional influences of LLPS in the context of select endogenous genes in live cells. Another key endeavor will be to decipher the molecular basis for the repressive effects of excessive LCD-LCD interactions on transcription even in the absence of LLPS. The quantitative single-cell imaging methods we developed to investigate EWS/FLI1 could pave the path to test the Goldilocks principle for other TFs in future studies.

**Fig. 5.**
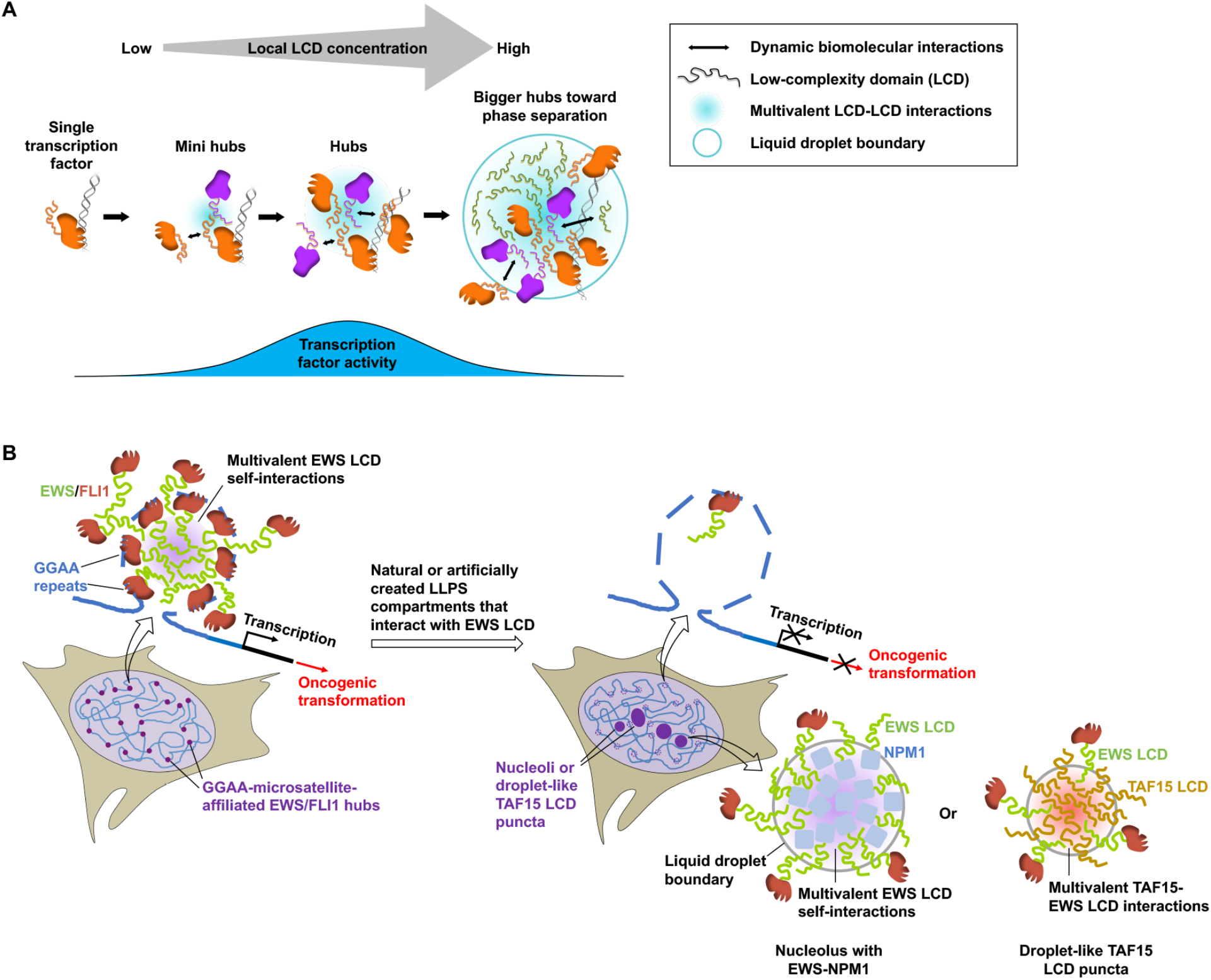
Models for LCD-interaction-mediated transcription. (**A**) Goldilocks principle of transcriptional activation: effective gene activation requires an optimal amount of multivalent LCD-LCD interactions. Overly high amounts of LCD-LCD interactions and LLPS at target genes repress transcription. (**B**) LLPS compartments formed away from EWS/FLI1 target genes (nucleoli containing EWS-NPM1 or droplet-like TAF15 LCD puncta) trap endogenous EWS/FLI1 within the compartments via homotypic or heterotypic LCD-LCD interactions, causing repression of EWS/FLI1-driven oncogenic transcription.

In the second part of this work, we demonstrated that multivalent EWS LCD self-interactions artificially created in the nucleolus are strong enough to relocate EWS/FLI1 from chromatin to the nucleolus, repress EWS/FLI1-driven transcription, and inhibit malignant transformation of Ewing’s sarcoma cells (Fig. 5B). Our findings reveal an important property of EWS LCD self-interactions, i.e., when recruited into the nucleolus, they can overcome EWS/FLI1-DNA binding interactions and effectively sequester TFs in a non-productive compartment. We expect similar strategies may allow trapping of EWS/FLI1 in other ectopic subcellular locations as well. The ability of subcellular mislocalization of LCD-LCD interactions to disrupt EWS/FLI1-chromatin binding and transactivation suggests a potentially novel therapeutic strategy for Ewing’s sarcoma. If the EWS-NPM1 protein we used could be replaced by smaller and more deliverable drug-like molecules that specifically interact with the EWS LCD, one could imagine using drug-induced mislocalization or sequestration of the target protein to an ectopic subcellular structure to disrupt its function. Moreover, similar approaches can potentially be used to target many other disease-causing TFs, which are notoriously difficult targets for the development of molecular therapies.

Our work on nucleolar targeting additionally sheds light on methods to more rigorously diagnose LLPS in vivo. Differential molecular diffusion dynamics provides a rigorous but seldom applied criterion for defining LLPS in vivo. In previous studies where molecular diffusion was measured by spaSPT, the lack of differences in diffusion dynamics inside versus outside a presumed LLPS compartment revealed that the compartment was actually not formed by LLPS *(51, 52)*. Here, by measuring diffusion of EWS/FLI1 in the nucleolus versus the nucleoplasm, we found that it diffuses much more slowly in the nucleolus, as expected of this well-studied, bone fide LLPS compartment. This observation provides a proof of concept that spaSPT measurements of protein diffusion rates can serve as an effective means of detecting LLPS based on differential viscosities within live cells. However, there remain clear limitations in our current method for single-molecule diffusion measurements, especially within submicron-sized puncta. Indeed, we were unable to precisely quantify diffusion rates within TAF15 LCD puncta that are significantly smaller than nucleoli and that often fall in the submicron range. Thus, here we observed apparent LLPS of TAF15 LCD based on the prominent droplet-like behaviors of TAF15 LCD puncta, but refrain from concluding that the puncta are true LLPS condensates.

It seems clear, however, that TAF15 LCD puncta colocalize with EWS/FLI1 and markedly repress EWS/FLI1-driven transcription. Besides causing a redistribution of EWS/FLI1 and transcription machinery as discussed above (Fig. 5B), another functionally relevant consequence of apparent LLPS might be that droplet-like TAF15 LCD puncta confine diffusion of transcription regulators and change their diffusion rates within the puncta. It remains unknown how such changes of protein dynamics at genes affect transactivation, and future studies are required to reveal potential functional consequences. While we have studied effects on specific endogenous target genes of EWS/FLI1, it will be important in the future to investigate how tuning the level of LCD-LCD interactions to form larger hubs and LLPS affects the global transcriptome of a cell.

## Acknowledgments

We thank L. Lavis for providing fluorescent HaloTag ligands; Q. Zhu for help with molecular cloning; K. Adiga for help optimizing conditions of doxycycline induction; the CRL Flow Cytometry Facility and Molecular Imaging Center; A. Hansen, S. Teves, and members of Tjian/Darzacq labs for critical reading of the manuscript.

## Author contributions

Conceptualization: RT, SC

Funding acquisition: RT, XD, CDD

Investigation: SC, TGWG, CDD, GMD, RT

Software: SC, TGWG

Visualization: SC, TGWG, CDD

Supervision: RT

Project administration: RT, SC

Writing – original draft: SC, RT, TGWG, CDD

Writing – review & editing: SC, RT, TGWG, CDD, XD

## References

1. J. T. Kadonaga, A. J. Courey, J. Ladika, R. Tjian, Distinct regions of Sp1 modulate DNA binding and transcriptional activation. Science. 242, 1566–1570 (1988).

2. S. Oka et al., NMR structure of transcription factor Sp1 DNA binding domain. Biochemistry. 43, 16027–16035 (2004).

3. Y. E. Guo et al., Pol II phosphorylation regulates a switch between transcriptional and splicing condensates. Nature. 572, 543–548 (2019).

4. M. Boehning et al., RNA polymerase II clustering through carboxy-terminal domain phase separation. Nat. Struct. Mol. Biol. 25, 833–840 (2018).

5. I. Kwon et al., Phosphorylation-regulated binding of RNA polymerase II to fibrous polymers of low-complexity domains. Cell. 155, 1049–1060 (2013).

6. B. R. Sabari et al., Coactivator condensation at super-enhancers links phase separation and gene control. Science. 361, 10.1126/science.aar3958. Epub 2018 Jun 21 (2018).

7. W. K. Cho et al., Mediator and RNA polymerase II clusters associate in transcription-dependent condensates. Science. 361, 412–415 (2018).

8. S. Chong et al., Imaging dynamic and selective low-complexity domain interactions that control gene transcription. Science. 361, 10.1126/science.aar2555. Epub 2018 Jun 21 (2018).

9. M. Mir et al., Dynamic multifactor hubs interact transiently with sites of active transcription in Drosophila embryos. Elife. 7, 10.7554/eLife.40497 (2018).

10. M. Mir et al., Dense Bicoid hubs accentuate binding along the morphogen gradient. Genes Dev. 31, 1784–1794 (2017).

11. J. Dufourt et al., Temporal control of gene expression by the pioneer factor Zelda through transient interactions in hubs. Nat. Commun. 9, 5194–z (2018).

12. D. T. McSwiggen, M. Mir, X. Darzacq, R. Tjian, Evaluating phase separation in live cells: diagnosis, caveats, and functional consequences. Genes Dev. 33, 1619–1634 (2019).

13. D. Hnisz, K. Shrinivas, R. A. Young, A. K. Chakraborty, P. A. Sharp, A Phase Separation Model for Transcriptional Control. Cell. 169, 13–23 (2017).

14. J. E. Henninger et al., RNA-Mediated Feedback Control of Transcriptional Condensates. Cell. 184, 207–225.e24 (2021).

15. A. Boija et al., Transcription Factors Activate Genes through the Phase-Separation Capacity of Their Activation Domains. Cell. 175, 1842–1855.e16 (2018).

16. M. T. Wei et al., Nucleated transcriptional condensates amplify gene expression. Nat. Cell Biol. 22, 1187–1196 (2020).

17. N. Schneider et al., Liquid-liquid phase separation of light-inducible transcription factors increases transcription activation in mammalian cells and mice. Sci. Adv. 7, 10.1126/sciadv.abd3568. Print 2021 Jan (2021).

18. J. Trojanowski, L. Frank, A. Rademacher, P. Grigaitis, K. Rippe, Transcription activation is enhanced by multivalent interactions independent of phase separation. bioRxiv., 2021.01.27.428421 (2021).

19. J. A. Riback et al., Composition-dependent thermodynamics of intracellular phase separation. Nature. 581, 209–214 (2020).

20. C. P. Brangwynne, T. J. Mitchison, A. A. Hyman, Active liquid-like behavior of nucleoli determines their size and shape in Xenopus laevis oocytes. Proc. Natl. Acad. Sci. U. S. A. 108, 4334–4339 (2011).

21. K. M. Johnson et al., Role for the EWS domain of EWS/FLI in binding GGAA-microsatellites required for Ewing sarcoma anchorage independent growth. Proc. Natl. Acad. Sci. U. S. A. 114, 9870–9875 (2017).

22. G. Boulay et al., Cancer-Specific Retargeting of BAF Complexes by a Prion-like Domain. Cell. 171, 163–178.e19 (2017).

23. L. Zuo et al., Loci-specific phase separation of FET fusion oncoproteins promotes gene transcription. Nat. Commun. 12, 1491–7 (2021).

24. N. Guillon et al., The oncogenic EWS-FLI1 protein binds in vivo GGAA microsatellite sequences with potential transcriptional activation function. PLoS One. 4, e4932 (2009).

25. K. Gangwal et al., Microsatellites as EWS/FLI response elements in Ewing’s sarcoma. Proc. Natl. Acad. Sci. U. S. A. 105, 10149–10154 (2008).

26. T. L. Lenstra, J. Rodriguez, H. Chen, D. R. Larson, Transcription Dynamics in Living Cells. Annu. Rev. Biophys. 45, 25–47 (2016).

27. M. Altmeyer et al., Liquid demixing of intrinsically disordered proteins is seeded by poly(ADP-ribose). Nat. Commun. 6, 8088 (2015).

28. K. M. Johnson, C. Taslim, R. S. Saund, S. L. Lessnick, Identification of two types of GGAA-microsatellites and their roles in EWS/FLI binding and gene regulation in Ewing sarcoma. PLoS One. 12, e0186275 (2017).

29. D. M. Mitrea et al., Nucleophosmin integrates within the nucleolus via multi-modal interactions with proteins displaying R-rich linear motifs and rRNA. Elife. 5, 10.7554/eLife.13571 (2016).

30. D. M. Mitrea et al., Self-interaction of NPM1 modulates multiple mechanisms of liquid-liquid phase separation. Nat. Commun. 9, 842–3 (2018).

31. F. Witzel, R. Fritsche-Guenther, N. Lehmann, A. Sieber, N. Bluthgen, Analysis of impedance-based cellular growth assays. Bioinformatics. 31, 2705–2712 (2015).

32. S. L. Lessnick, C. S. Dacwag, T. R. Golub, The Ewing’s sarcoma oncoprotein EWS/FLI induces a p53-dependent growth arrest in primary human fibroblasts. Cancer. Cell. 1, 393–401 (2002).

33. B. L. Sprague, R. L. Pego, D. A. Stavreva, J. G. McNally, Analysis of binding reactions by fluorescence recovery after photobleaching. Biophys. J. 86, 3473–3495 (2004).

34. A. S. Hansen, I. Pustova, C. Cattoglio, R. Tjian, X. Darzacq, CTCF and cohesin regulate chromatin loop stability with distinct dynamics. Elife. 6, 10.7554/eLife.25776 (2017).

35. J. Elf, G. W. Li, X. S. Xie, Probing transcription factor dynamics at the single-molecule level in a living cell. Science. 316, 1191–1194 (2007).

36. A. S. Hansen et al., Robust model-based analysis of single-particle tracking experiments with Spot-On. Elife. 7, 10.7554/eLife.33125 (2018).

37. J. B. Grimm et al., Bright photoactivatable fluorophores for single-molecule imaging. Nat. Methods. 13, 985–988 (2016).

38. L. Xiang, K. Chen, R. Yan, W. Li, K. Xu, Single-molecule displacement mapping unveils nanoscale heterogeneities in intracellular diffusivity. Nat. Methods. 17, 524–530 (2020).

39. A. Heckert, L. Dahal, R. Tjian, X. Darzacq, Recovering mixtures of fast diffusing states from short single particle trajectories. bioRxiv., 2021.05.03.442482 (2021).

40. W. S. Dynan, R. Tjian, The promoter-specific transcription factor Sp1 binds to upstream sequences in the SV40 early promoter. Cell. 35, 79–87 (1983).

41. Y. Shin et al., Liquid Nuclear Condensates Mechanically Sense and Restructure the Genome. Cell. 176, 1518 (2019).

42. Y. Shin et al., Spatiotemporal Control of Intracellular Phase Transitions Using Light-Activated optoDroplets. Cell. 168, 159–171.e14 (2017).

43. D. Bracha et al., Mapping Local and Global Liquid Phase Behavior in Living Cells Using Photo-Oligomerizable Seeds. Cell. 176, 407 (2019).

44. J. W. Gibbs, On the Equilibrium of Heterogeneous Substances (Transactions of the Connecticut Academy of Arts and Sciences. 1876), pp. 108–248.

45. Thomas Graham, X. Liquid diffusion applied to analysis. Philosophical Transactions of the Royal Society. 151, 183–224 (1861).

46. S. Chong, M. Mir, Towards Decoding the Sequence-Based Grammar Governing the Functions of Intrinsically Disordered Protein Regions. J. Mol. Biol. 433, 166724 (2021).

47. G. J. Narlikar et al., Is transcriptional regulation just going through a phase? Mol. Cell. 81, 1579–1585 (2021).

48. S. Boeynaems et al., Protein Phase Separation: A New Phase in Cell Biology. Trends Cell Biol. 28, 420–435 (2018).

49. Y. Shin, C. P. Brangwynne, Liquid phase condensation in cell physiology and disease. Science. 357, 10.1126/science.aaf4382 (2017).

50. S. F. Banani, H. O. Lee, A. A. Hyman, M. K. Rosen, Biomolecular condensates: organizers of cellular biochemistry. Nat. Rev. Mol. Cell Biol. 18, 285–298 (2017).

51. S. Collombet et al., RNA polymerase II depletion from the inactive X chromosome territory is not mediated by physical compartmentalization. bioRxiv., 2021.03.26.437188 (2021).

52. D. T. McSwiggen et al., Evidence for DNA-mediated nuclear compartmentalization distinct from phase separation. Elife. 8, 10.7554/eLife.47098 (2019).

