## Supplementary materials for "Tuning levels of low-complexity domain interactions to modulate endogenous oncogenic transcription"

**This PDF file includes:**

Materials and Methods
Figs. S1 to S11
References (*53-55)*

### Materials and Methods

Cell culture and stable cell line construction

The knock-in A673 cell line expressing endogenous EWS/FLI1-Halo *(8)* were grown in high-glucose DMEM (ThermoFisher, 10566016) with 10% FBS and 1% penicillin-streptomycin. For live-cell imaging, the medium was identical except that phenol-red-free DMEM (ThermoFisher, 31053028) was used. The knock-in cells and those with modifications described below were cultured at 37°C with 5% $CO_2$.

We engineered the knock-in line to express high levels of mNG-EWS-NPM1 or mNG-NPM1 upon doxycycline induction using PiggyBac transposition, drug selection, and clone selection. First, we cloned the cDNA encoding mNG-EWS-NPM1 or mNG-NPM1 into an Xlone vector *(53)* containing PiggyBac elements, a Tet-On 3G inducible gene expression system, and a puromycin resistance gene. We then co-transfected the knock-in cells with the Xlone vector and a SuperPiggyBac transposase vector using Lipofectamine 3000 transfection reagent (ThermoFisher Scientific, L300015) following the manufacturer's instructions. 72 hours after transfection, we started with selection by adding 1 μg/ml puromycin. Untransfected knock-in cells were treated with puromycin in parallel, and selection was judged to be complete once no untransfected cells remained (~ 3 days). To make a clonal cell line with sufficiently high induction of mNG-EWS-NPM1 to sequester endogenous EWS/FLI1-Halo to the nucleolus, we plated the puromycin-selected cells one cell per well into 96-well plates by limiting dilution, expanded single clones, and screened for clones by fluorescence imaging with a high-throughput automated confocal microscope (Perkin Elmer, Opera Phenix, 40x water objective). To prepare for live-cell imaging samples, we plated clones in 96-well plates (Perkin Elmer, CellCarrier-96), induce expression of mNG-EWS-NPM1 with doxycycline (100~200 ng/ml) for 96 hours, stain cells with 200 nM JF646 HaloTag ligand for 15 min, and washed twice (each wash: remove medium, rinse twice with PBS, incubate in fresh medium for 30 min). At the end of the final wash, the medium was changed to phenol-red-free imaging medium as described above. After identifying a desirable clonal cell line with high induction of mNG-EWS-NPM1, we isolated single clones of puromycin-selected cells with inducible expression of mNG-NPM1 and used a flow cytometer (BD Biosciences, BD LSRFortessa Cell Analyzer) to screen for clones with mNG-NPM1 levels similar to mNG-EWS-NPM1 as described above under the same induction condition.

Fluorescence in situ hybridization (FISH)

The knock-in A673 cells were plated on 18 mm circular No. 1 cover glasses (VWR VistaVision, 16004-300) and transfected with a protein expression plasmid using Lipofectamine 3000. 24 hours after transfection, we stained the cells with 200 nM JFX549 HaloTag ligand following the protocol described above, fixed the cells, and then proceeded with FISH. To measure nascent transcription levels of *ABHD6*, *CAV1*, and *GAPDH* genes, we performed intron RNA FISH following the published Stellaris RNA FISH protocol for adherent cells (https://biosearchassets.blob.core.windows.net/assets/bti_stellaris_protocol_adherent_cell.pdf) using Quasar 670-labeled FISH probes designed with the online software Stellaris Probe Designer (https://www.biosearchtech.com/support/tools/design-software/stellaris-probe-designer) and purchased from LGC Biosearch Technologies. To visualize *ABHD6* and *CAV1* gene loci with DNA FISH, we prepared FISH probes targeting each gene following the procedure described in *(8)* and performed 3D DNA FISH following the published protocol *(54)*.

Confocal fluorescence imaging and analyses for protein and nucleic acid distribution

Two confocal microscopes were used to image intron RNA FISH samples. One is an inverted laser scanning confocal microscope (Zeiss, LSM 710 AxioObserver) equipped with 34-channel spectral detection, a motorized stage, a full incubation chamber maintaining 37°C and 5% $CO_2$, a heated stage, an X-Cite 120 illumination source as well as several laser lines (405, 458, 488, 514, 561, 591, 633 nm). Images were acquired with a 40x Plan NeoFluar NA1.3 oil-immersion objective under control of the Zeiss Zen software. The other is an inverted laser scanning confocal microscope with Airyscan super-resolution capability (Zeiss, LSM 900 with Airyscan 2) and equipped with four laser lines (405, 488, 561, 640 nm). Images were acquired with a 40x oil objective (Zeiss Plan-Apochromat 40x/1.3 Oil DIC) in the confocal (CO) mode under control of the Zen software. We acquired z stacks of RNA FISH samples with a slice interval of 0.3 μm. 405 nm, 488 nm, 561 nm, and 633 or 640 nm lasers were used to excite the fluorescence of Hoechst-labeled nuclei, EGFP or mNG-labeled proteins, JFX549-labeled EWS/FLI1-Halo, and Quasar 670-labeled intron RNA FISH, respectively. Before acquiring any fluorescence image, we carefully set the laser intensity and microscope detectors to make sure that no pixel in the image was saturated. We used proper emission filters for sequential four-color imaging and ensured no bleed-through between the four channels by imaging cell samples that contain only one of the four fluorophores (Hoechst, EGFP or mNG, JFX549, and Quasar 670) under the four-color imaging settings. To quantify intron RNA FISH intensities, we made z projection for each z stack that results in a sum intensity image, integrated the intensity of all pixels within a gene locus on the sum image, and subtracted the product of the average intranuclear background intensity and the pixel number within the gene locus. In box plots of intron RNA FISH intensities (Fig. 1D-F, 2E-G, 3C-E, S1B-C, S3B-C, & S5B-E), corresponding fluorescent protein intensities were the average intranuclear intensities on the sum images generated from z projection of the protein z-stack images.

DNA FISH samples were imaged on the confocal microscope (Zeiss, LSM 900 with Airyscan 2) in the super-resolution (SR) mode using the 40x oil objective described above. We acquired z stacks of DNA FISH samples with a slice interval of 0.2 μm. 405 nm, 488 nm, and 640 nm lasers were used to excite the fluorescence of Hoechst-labeled nuclei, EGFP-TAF15, and Cy5-labeled DNA FISH, respectively. We made sure no pixel in the image was saturated and there was no bleed-through between three channels using the approaches described above.

For live-cell imaging of fluorescent protein distribution (Fig. 1A, 2A, & 3A), the knock-in A673 cells were grown on glass-bottom (No. 1.5, 14 mm diameter) 35 mm dishes (MatTek, P35G-1.5-14-C), transfected with a protein expression plasmid using Lipofectamine 3000, and stained the cells with 200 nM JF646 or JFX549 HaloTag ligand. The live-cell samples were imaged on the Airyscan confocal microscope in the super-resolution (SR) mode using the 40x oil objective. We acquired z stacks of the samples with a slice interval of 0.2 μm. We made sure no pixel in the image was saturated and there was no bleed-through between channels. Protein enrichment at the endogenous EWS/FLI1-Halo hubs (Fig. 1B, 2B, & S6) was examined by image analysis methods described in *(8)*. Radial intensity profiles were plotted using a published ImageJ plugin “Radial Profile Plot” (https://imagej.nih.gov/ij/plugins/radial-profile.html).

Fluorescence recovery after photobleaching (FRAP)

FRAP was performed on the inverted laser scanning confocal microscope (Zeiss, LSM 710 AxioObserver) described above. The 561 nm laser and the epi-illumination mode were used for FRAP measurements. Images were acquired with a 40x Plan NeoFluar NA1.3 oil-immersion objective. The knock-in A673 cells were grown on glass-bottom (No. 1.5, 14 mm diameter) 35 mm dishes (MatTek, P35G-1.5-14-C). To measure the FRAP dynamics of EWS/FLI1-Halo in the nucleolus, we transfected the knock-in cells with a plasmid encoding mNG-EWS-NPM1 and stained the cells with 500 nM HaloTag TMR ligand (Promega, G8251) following the protocol described above. We acquired 1000 frames at one frame per 0.3 seconds with the first 5 frames acquired before the bleach pulse for the measurement of baseline fluorescence of the bleach spot and the whole nucleus. We chose to photobleach a circular spot (radius ~1 µm) within a nucleolus using the 561 nm laser at maximum intensity. We extracted FRAP curves from the acquired movies using methods described in *(8)*. To measure the FRAP dynamics of EWS/FLI1-Halo in the nucleoplasm, we followed the same procedure as above, except that the knock-in cells were not transfected and a circular bleach spot (radius ~ 1 µm) was chosen within the nucleoplasm of a cell and at least 1 µm from nuclear and nucleolar boundaries.

Stroboscopic photo-activatable single particle tracking (spaSPT) and analyses

The knock-in cells with inducible expression of mNG-EWS-NPM1 were grown on 25 mm circular No. 1.5 cover glasses (Azer Scientific, 200251) that were plasma-cleaned prior to use. We induced the cells with 200 ng/ml of doxycycline for 96 hours, stained the cells with 20 nM PA-JF646 and 200 nM JFX549 HaloTag ligands, and performed single-molecule imaging of EWS/FLI1-Halo on a custom-built Nikon (Nikon Instruments Inc.) TI microscope described in *(34)*. We took images with a 100x/NA 1.49 oil-immersion TIRF objective (Nikon apochromat CFI Apo TIRF 100x Oil) under highly inclined and laminated optical sheet (HILO) illumination *(55)* using following laser lines: 488 nm for mNG; 561 nm for JFX549; 405 nm and 633 nm for photo-activation and excitation of PA-JF646, respectively. The incubation chamber maintained a humidified 37°C atmosphere with 5% $CO_2$ and the objective was similarly heated to 37°C for live-cell experiments.

High-concentration JFX549 staining allows visualization of the intracellular distribution of EWS/FLI1-Halo. We chose cells with EWS/FLI1-Halo significantly enriched in the nucleolus to perform spaSPT (Fig. 4B). The procedure of spaSPT largely follows what is described in *(34)*. Both the excitation laser (633 nm) and the photo-activation laser (405 nm) for PA-JF646 were pulsed. Each frame consisted of a 7-ms camera exposure time followed by a ~500 µs camera 'dead' time. The excitation laser (633 nm) was pulsed for 1 ms starting at the beginning for the 7 ms camera exposure time. The photo-activation laser (405 nm) was pulsed during the ~500 µs camera 'dead' time, minimizing fluorescence background. Each cell was imaged for 20,000 frames corresponding to ~1.5 min. We recorded movies for 20 cells per day for 4 days. Over 400,000 single EWS/FLI1-Halo molecules were localized and tracked using the quot package (https://github.com/alecheckert/quot) *(39)*. All the downstream analysis was performed using custom MATLAB code.

To prepare binary masks of nuclei and nucleoli, images of mNG-EWS-NPM1 collected before and after each single-molecule movie were first averaged and then filtered using Gaussian filters of width 1.5 pixels for nucleoli and 5 pixels for nuclei. Filtered images were then binarized using the threshold

$$T = I_{min} + A(I_{max} - I_{min})$$

where $I_{min}$ and $I_{max}$ are the minimum and maximum pixel intensities and $A$ is a constant, which was set to 0.05 for nuclei and 0.3 for nucleoli. These binary masks were used to classify each single-molecule localization as nucleolar, nucleoplasmic, or cytoplasmic. Trajectories were classified as nucleolar or nucleoplasmic if all localizations in that trajectory fell within the nucleolus or nucleoplasm, respectively (Fig. 4C).

Sorted trajectories from multiple cells were pooled by subnuclear compartment and fit to a 2-state model using Spot-On *(36)* with the following parameters:

| **Parameter** | **Value** | **Description** |
|---|---|---|
| TimeGap | 7.5 | Delay between frames in ms |
| dZ | 0.7 | Depth of field in μm |
| GapsAllowed | 0 | Gaps not allowed in tracking |
| TimePoints | 8 | Consider 8 timepoints, i.e., 7 delays |
| BinWidth | 0.01 | Use bin width of 0.01 μm for displacement histogram |
| UseEntireTraj | 0 | Consider only 4 jumps for each delay from each trajectory |
| JumpsToConsider | 4 | |
| MaxJump | 5.05 | Maximal displacement to consider in μm |
| ModelFit | 2 | CDF fitting |
| DoSingleCellFit | 0 | Do not fit data for each cell individually |
| NumberOfStates | 2 | Use 2-state model |
| FitIterations | 2 | Number of fitting iterations |
| FitLocError | 1 | Fit localization error from the data |
| FitLocErrorRange | [0.01, 0.075] | Minimum and maximum allowable localization error |
| UseWeights | 0 | Weight all time points equally |
| D_Free_2State | [0.2 25] | Minimum and maximum allowable diffusion coefficient for "free" population |
| D_Bound_2State | [0.0001 0.05] | Minimum and maximum allowable diffusion coefficient for "bound" population |

Bootstrap confidence intervals for fit parameters were determined by randomly resampling data by cell with replacement. Spot-On was used to analyze 200 bootstrap replicates, and a 95% confidence interval was estimated as the mean value of each parameter plus or minus 1.96 standard deviations.

Randomized nucleolar masks were prepared using a sequence of 10000 Monte Carlo moves (Fig. S10). At each step, each nucleolus (i.e., connected component in the nucleolar mask) was displaced horizontally by -1, 0, or +1 pixels, displaced vertically by -1, 0, or +1 pixels, and rotated by -10°, 0°, or +10°. Each displacement or rotation was set to occur with an equal probability. Alternatively, "teleportation" to a new random location within the nuclear mask was performed with a probability of 1% per step. This new configuration was accepted if all nucleoli remained within the nucleus and did not overlap, and it was rejected otherwise. The current configuration was written out every 100 steps. Spot-On was used to analyze a total of 200 randomized mask replicates. For each replicate, one randomized mask was chosen per cell from

Monte Carlo steps 1000 and above. Sorting of nucleolar and nucleoplasmic trajectories and analysis using Spot-On was performed as described above.

RT-qPCR and analyses

Total RNA was purified from cells (treated or not with 150 ng/mL of doxycycline for 96 hours) using Trizol (ThermoFisher Scientific, 15596026) and quantified by Nanodrop. 500 ng of total RNA was retrotranscribed to cDNA using Maxima First Strand cDNA Synthesis Kit for RT-qPCR, with dsDNase (ThermoFisher Scientific, K1671). 5 µl of 1:20 cDNA dilutions were used for quantitative PCR with SYBR Select Master Mix for CFX (Applied Biosystems, ThermoFisher) on a BIO-RAD CFX Real-time PCR system. Two biological replicates and three technical replicates for each were performed for each gene. To normalize the data, we used the average value of 4 invariant transcripts (GAPDH, MED12, NAE1, BGALT3) *(8)* (Fig. S8) and calculated the normalized fold change for each target gene (Fig. 3F). Primers used were as described in *(8)*.

xCELLigence assay

The Cell Index (a representation of cell growth and viability) was measured in real time using the xCELLigence real-time cell analysis (RTCA) single plate instrument (RTCA-SP) (Acea Biosciences) following manufacturer's instructions. Cells were seeded at a density of 4000 cells/well in an RTCA E-plate View 96 (Agilent, 300601010). After 24 hours, protein expression (mNG-NPM1 or mNG-EWS-NPM1) was induced by replacing the media with fresh media containing 150 ng/mL of doxycycline. Media was changed every 48 days afterwards. The Cell Index (graphed as Cell Number in Fig. S9) was normalized to the value measured 7 hours after induction for each condition.

Soft agar colony formation assay

To examine how induction of mNG-EWS-NPM1 or mNG-NPM1 affects the malignant transformation phenotype of the knock-in A673 cells, we first seeded cells at a density of 6 x $10^4$ per well in a 6-well plate in 0.4% SeaPlaque GTG agarose (Fisher, BMA50111) and IMDM (ThermoFisher, 12200036) medium containing 20% FBS and 1% penicillin-streptomycin, added 0.3 ml liquid media (IMDM with 10% FBS and 1% penicillin-streptomycin) per well on top of agar, and cultured the cells in agar at 37°C with 5% $CO_2$ for 5 days. Then we induced protein expression by replacing the liquid media with new medium containing doxycycline that corresponds to 25 ng/ml in the entire cell culture volume (agar and liquid). We cultured the cells at 37°C with 5% $CO_2$ for another 10 days, changing doxycycline-containing liquid media every 3 days, and then stained the cells by applying to each well 200 µl of 1 mg/ml nitro blue tetrazolium chloride (NBT, ThermoFisher, N6495) in PBS. After culturing the stained cells in agar for 24 hours, we took images of the wells with a ChemiDoc MP imaging system (Bio-Rad). The live colonies appeared as distinct dots on agar (Fig. 3G). We counted the number of sizable colonies (diameter > 220 µm) for quantitative analyses. Inducible knock-in cells were cultured in agar without doxycycline induction in parallel. Two biological replicates and three technical replicates for each were performed for each condition. The average number of induced colonies is divided by the average number of uninduced colonies to calculate the fold change by protein induction. The fold changes by induction of mNG-EWS-NPM1 and mNG-NPM1 are plotted in Fig. 3H.

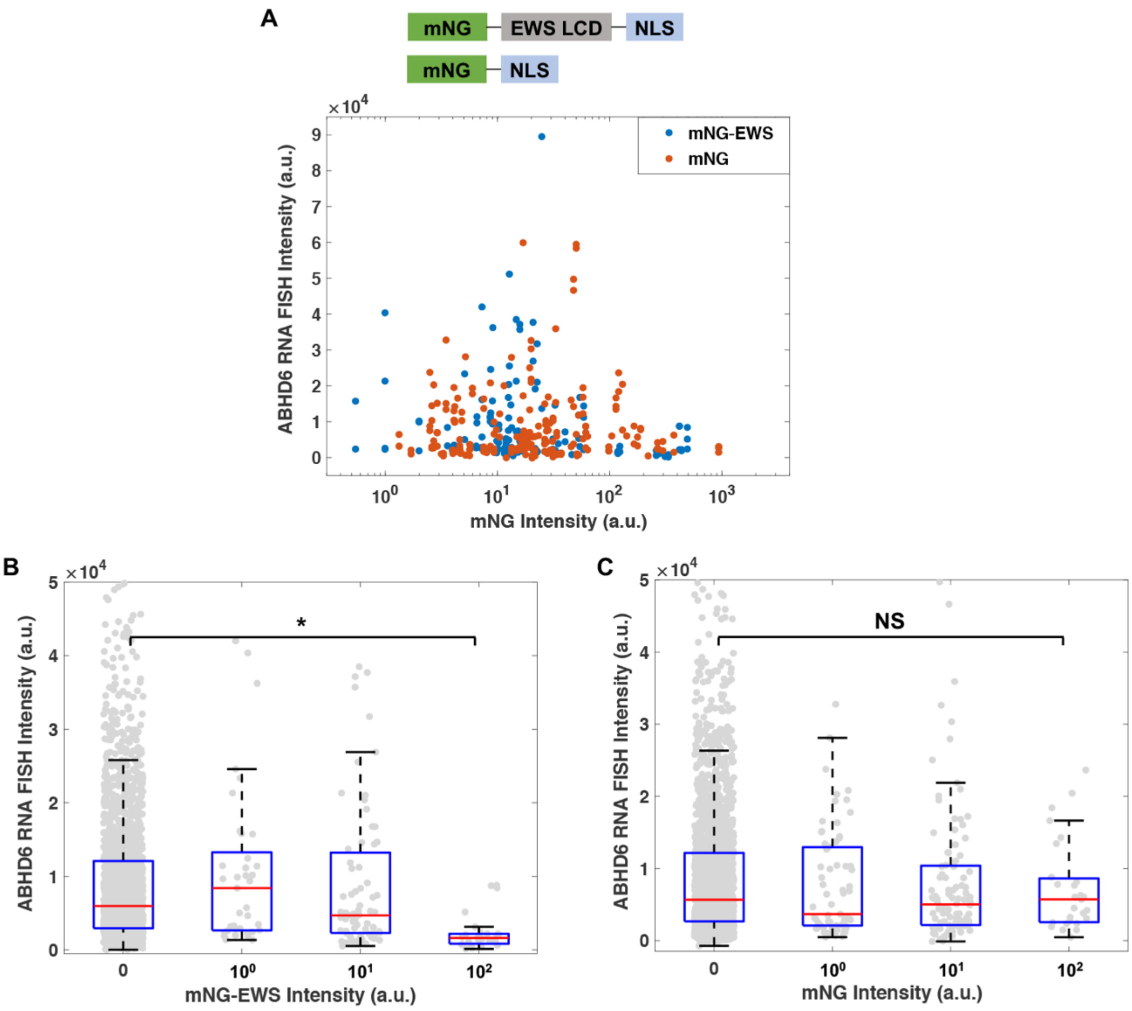


**Fig. S1. Expression of mNG-EWS but not mNG alone represses EWS/FLI1-driven transcription.**

(**A**) Intron RNA FISH intensities of individual *ABHD6* gene loci in the knock-in A673 cells expressing mNG-EWS (blue) or mNG alone (orange) tagged with a nuclear localization signal (NLS). Each dot represents one gene locus.
(**B-C**) Box plots of intron RNA FISH intensities of *ABHD6* gene after all the gene loci are categorized based on the corresponding nuclear mNG-EWS (**B**) or mNG (**C**) intensity. The x axis lists the order of magnitude of mNG-EWS (**B**) or mNG (**C**) intensities in each category. Individual gene loci are plotted in gray. mNG-negative cells after transfection (the “0” intensity category) are included here but not in (**A**). *: statistically significant decrease in the RNA FISH intensity of an mNG-positive category compared with an mNG-negative category ($p < 0.05$, two-sample t-test). NS: non-significant difference between two categories.

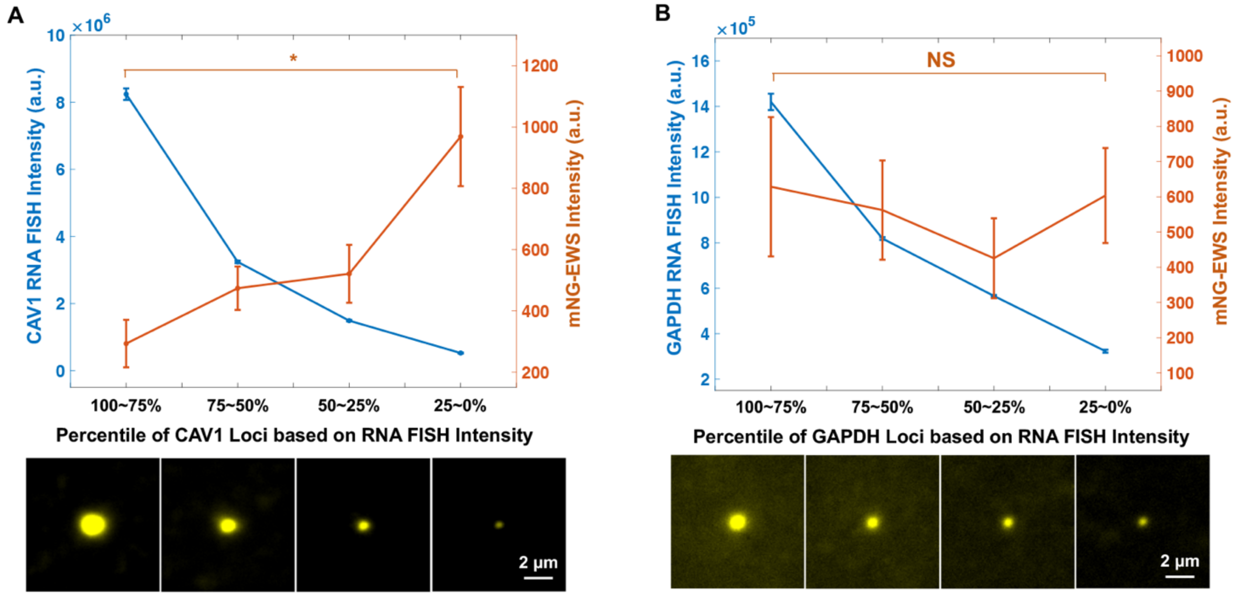


**Fig. S2. Inverse correlation between EWS LCD expression and EWS/FLI1-driven transcription.**

(**A**) (Upper) Averaged fluorescence intensities of intron RNA FISH (blue) and mNG-EWS (orange) at the *CAV1* gene loci within each percentile-based category. *: statistically significant increase in the mNG intensities of low-percentile *CAV1* gene loci based on the RNA FISH intensity compared with high-percentile loci ($p < 0.05$, two-sample t-test). Error bars represent bootstrapped standard deviation. (Bottom) Averaged intron RNA FISH images of *CAV1* gene loci in four categories based on the percentile rank of intron RNA FISH intensity. Each image is averaged from 211 gene loci.

(**B**) (Upper) Averaged fluorescence intensities of intron RNA FISH (blue) and mNG-EWS (orange) at the *GAPDH* gene loci within each percentile-based category. NS: non-significant difference between the mNG intensities of low-percentile and high-percentile *GAPDH* gene loci based on the RNA FISH intensity. (Bottom) Averaged intron RNA FISH images of *GAPDH* gene loci in four categories based on the percentile rank of intron RNA FISH intensity. Each image is averaged from 211 gene loci.

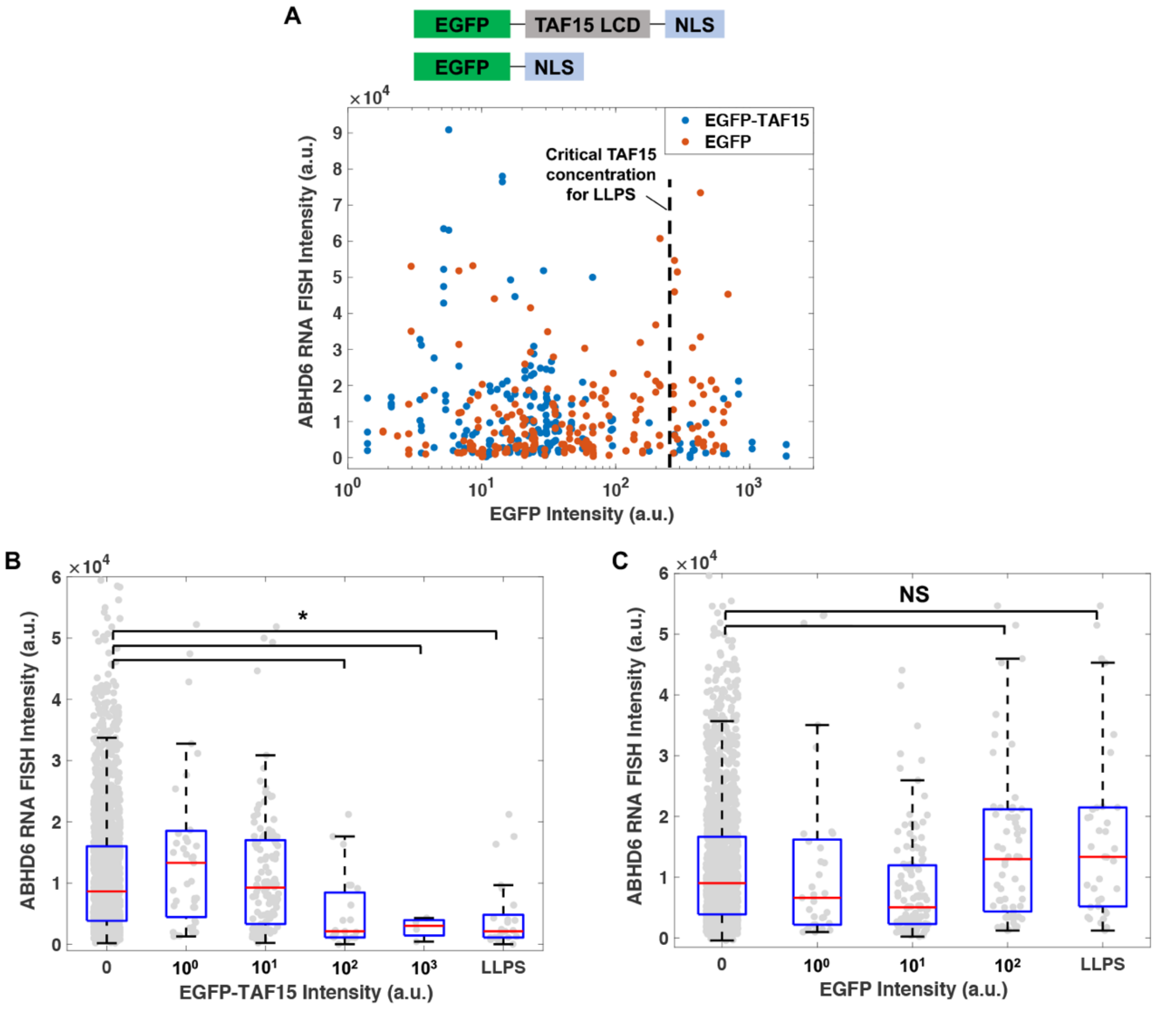


**Fig. S3. Expression of EGFP-TAF15 but not EGFP alone represses EWS/FLI1-driven transcription.**

(**A**) Intron RNA FISH intensities of individual *ABHD6* gene loci in the knock-in A673 cells expressing EGFP-TAF15 (blue) or EGFP alone (orange) tagged with an NLS. Each dot represents one gene locus. The dashed line marks the EGFP-TAF15 intensity above which apparent LLPS is detected. This intensity corresponds to the critical TAF15 concentration required for LLPS.

(**B-C**) Box plots of intron RNA FISH intensities of *ABHD6* gene after all the gene loci are categorized based on the corresponding nuclear EGFP-TAF15 (**B**) or EGFP (**C**) intensity. The x axis lists the order of magnitude of EGFP-TAF15 (**B**) or EGFP (**C**) intensities in each category. An additional "LLPS" category includes gene loci with EGFP-TAF15 (**B**) or EGFP (**C**) intensities above the critical value required for apparent LLPS of TAF15. Individual gene loci are plotted in gray. EGFP-negative cells after transfection (the "0" intensity category) are included here but not in (**A**). *: statistically significant decrease in the RNA FISH intensity of specific EGFP-positive categories compared with the EGFP-negative category ($p < 0.05$, two-sample t-test). NS: non-significant difference.

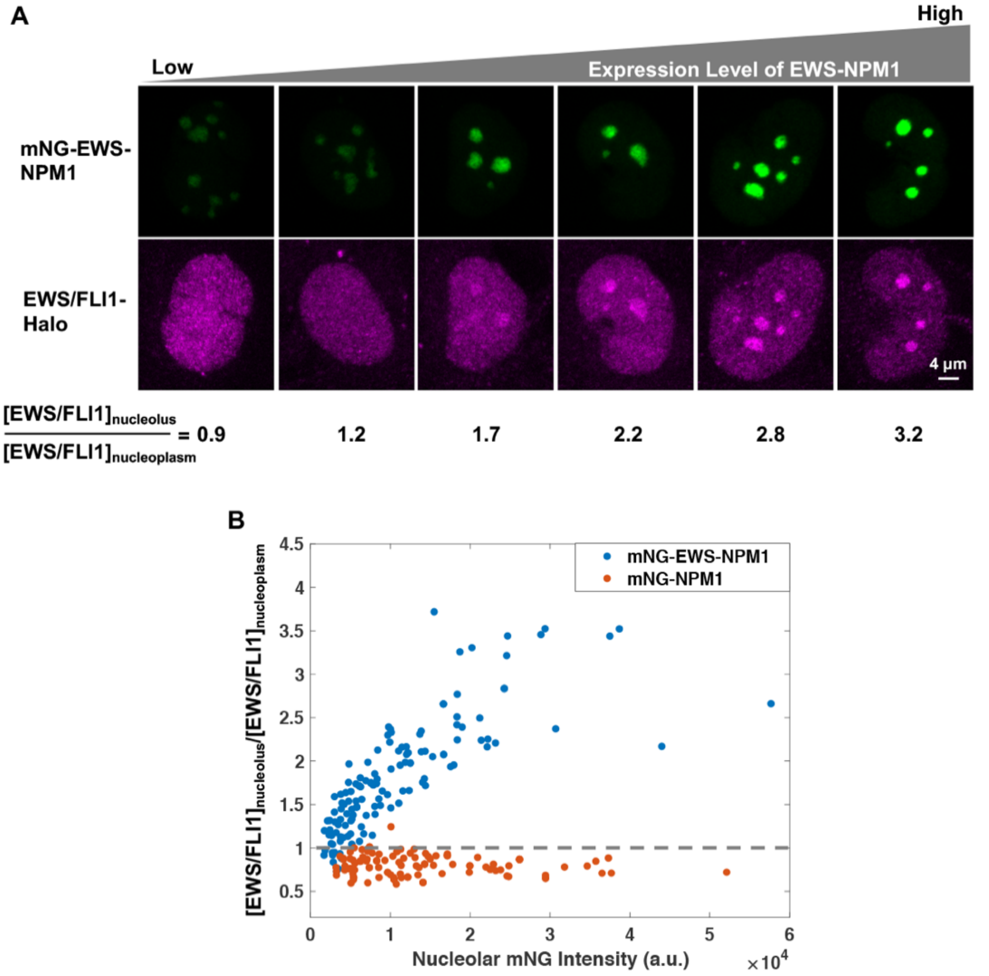


**Fig. S4. Overexpressed EWS-NPM1 but not NPM1 recruits endogenous EWS/FLI1 to the nucleolus.**

(**A**) Two-color images of single knock-in A673 cells with different expression levels of mNG-EWS-NPM1 (green). The nucleolar enrichment of endogenous EWS/FLI1-Halo (JFX549 labeled, magenta) increases with the expression level of EWS-NPM1, as quantified by the fluorescence intensity of mNG in the nucleolus. The nucleolar enrichment of EWS/FLI1 is quantified by the ratio of EWS/FLI1 concentration in the nucleolus to that in the nucleoplasm and listed below the respective cell. All single-cell images are acquired from the same sample with identical image acquisition settings and displayed with identical contrast settings.
(**B**) Quantified nucleolar enrichment of EWS/FLI1 ($[EWS/FLI1]_{nucleolus}/[EWS/FLI1]_{nucleoplasm}$ ratio) as a function of mNG-EWS-NPM1 (blue) or mNG-NPM1 (orange) expression level measured by the nucleolar mNG intensity. The ratio is above 1 for cells where EWS/FLI1 is recruited to the nucleolus.

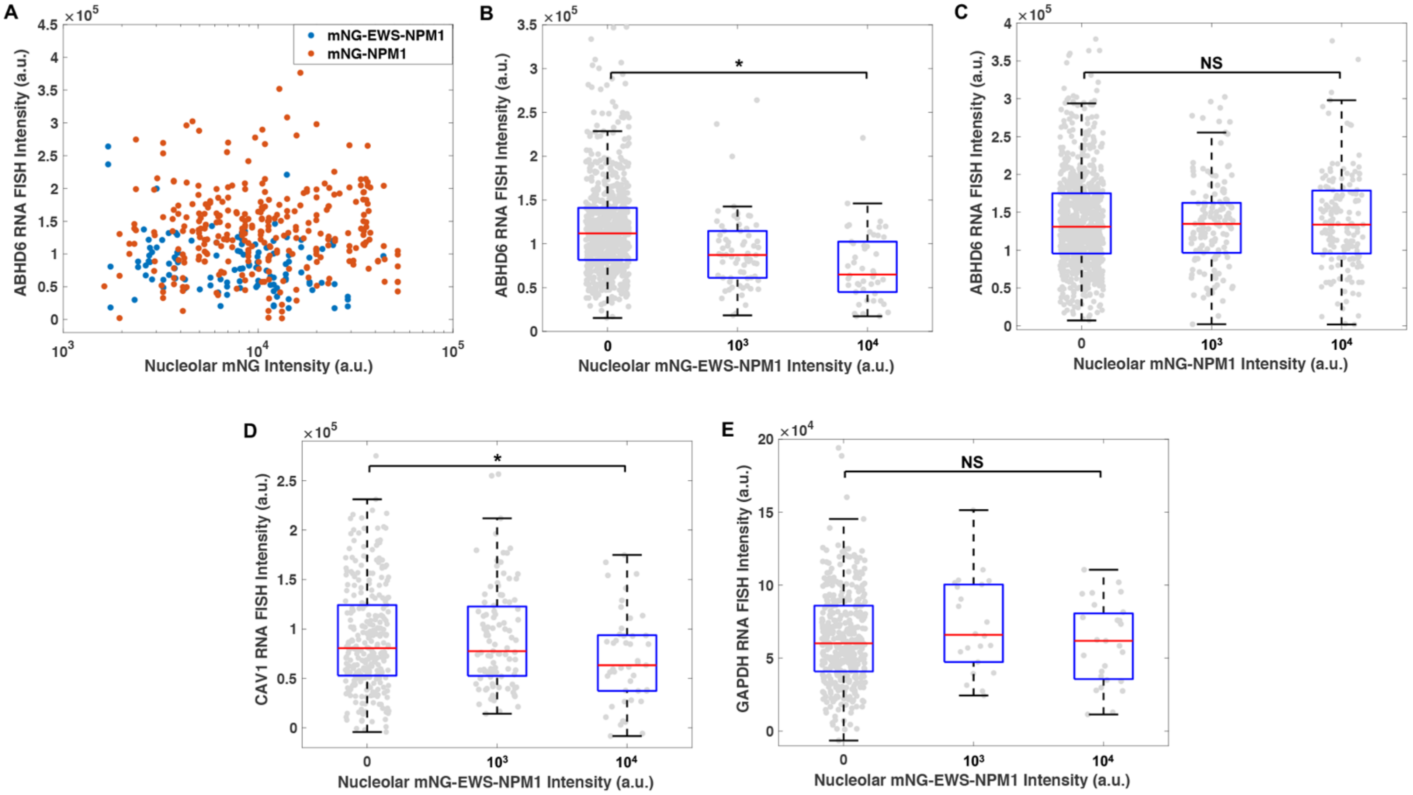


**Fig. S5. Overexpressed EWS-NPM1 but not NPM1 represses EWS/FLI1-driven transcription.**

(**A**) Intron RNA FISH intensities of individual *ABHD6* gene loci in the knock-in A673 cells expressing mNG-EWS-NPM1 (blue) or mNG-NPM1 (orange). Each dot represents one gene locus.

(**B-C**) Box plots of intron RNA FISH intensities of *ABHD6* gene after all the gene loci are categorized based on the corresponding nuclear mNG-EWS-NPM1 (**B**) or mNG-NPM1 (**C**) intensity. The x axis lists the order of magnitude of mNG-EWS-NPM1 (**B**) or mNG-NPM1 (**C**) intensities in each category. Individual gene loci are plotted in gray. mNG-negative cells after transfection (the "0" intensity category) are included here but not in (**A**). *: statistically significant decrease in the RNA FISH intensity of an mNG-positive category compared with an mNG-negative category ($p < 0.05$, two-sample t-test). NS: non-significant difference between two categories.

(**D-E**) Box plots of intron RNA FISH intensities of *CAV1* (**D**) or *GAPDH* (**E**) gene after all the gene loci are categorized based on the corresponding nuclear mNG-EWS-NPM1 intensity. Individual gene loci are plotted in gray. *: statistically significant decrease in the RNA FISH intensity of an mNG-positive category compared with an mNG-negative category ($p < 0.05$, two-sample t-test). NS: non-significant difference.

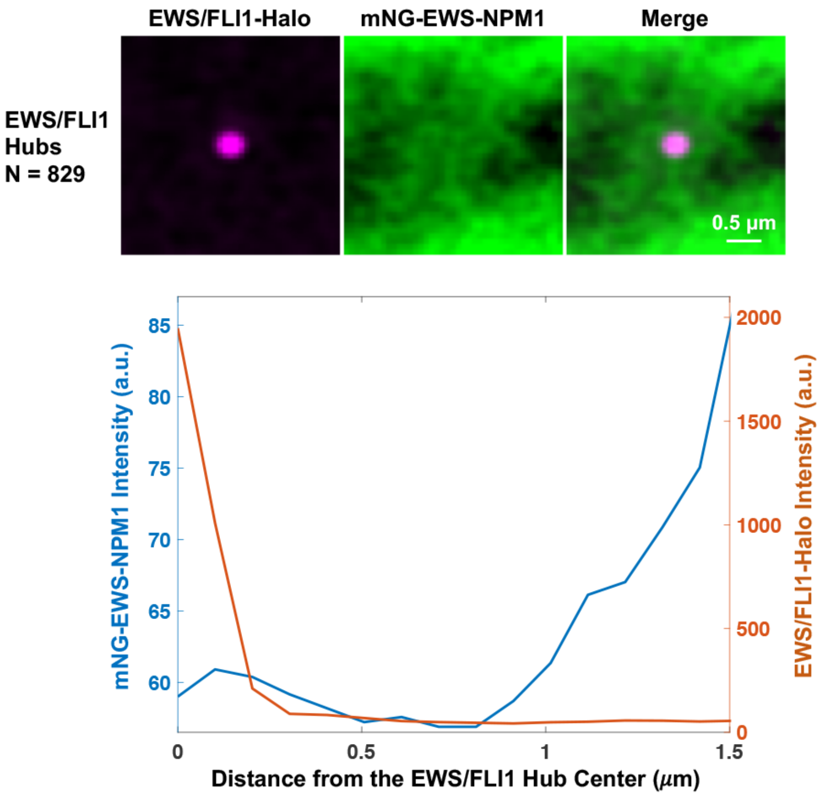


**Fig. S6. Overexpressed EWS-NPM1 does not accumulate at endogenous EWS/FLI1 hubs.**
(**A**) Averaged mNG-EWS-NPM1 (green) image at 829 EWS/FLI1 (magenta) hubs from 13 cells. Endogenously expressed EWS/FLI1-Halo is labeled with HaloTag ligand JF646.
(**B**) Radial intensity profiles of mNG-EWS-NPM1 (blue) and EWS/FLI1-Halo (orange) surrounding the center of EWS/FLI1 hubs on the averaged image.

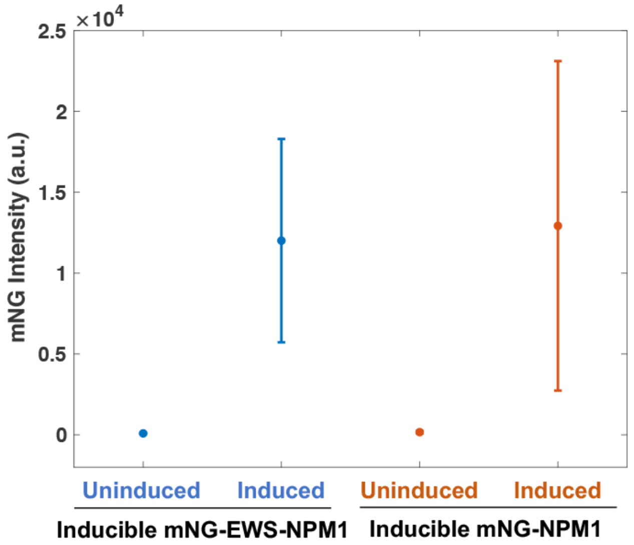


**Fig. S7. Two clonal cell lines based on the knock-in A673 line express similar levels of mNG-EWS-NPM1 and mNG-NPM1, respectively, upon doxycycline induction.**

Relative protein expression levels are estimated from mNG intensities of single living cells measured by flow cytometry, after the cells are induced by 150 ng/ml doxycycline for 96 hours. Error bars represent standard deviations.

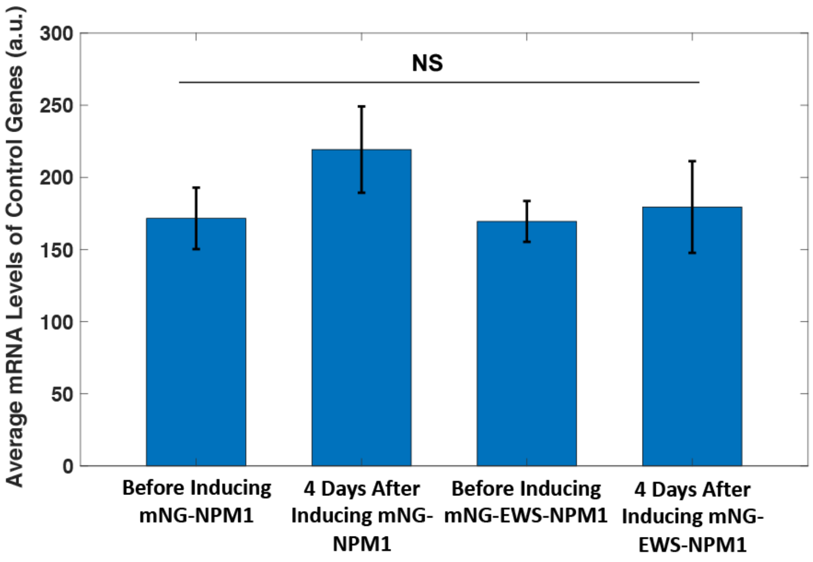


**Fig. S8. The average mRNA level of 4 non-EWS/FLI1-target genes (GAPDH, MED12, GALT3, and NAE1) in the knock-in A673 cells does not change by induction of EWS-NPM1 or NPM1 as measured using RT-qPCR.**

The average mRNA level of these invariant genes under each condition is used to normalize the mRNA level of each EWS/FLI1 target gene measured under the same condition.

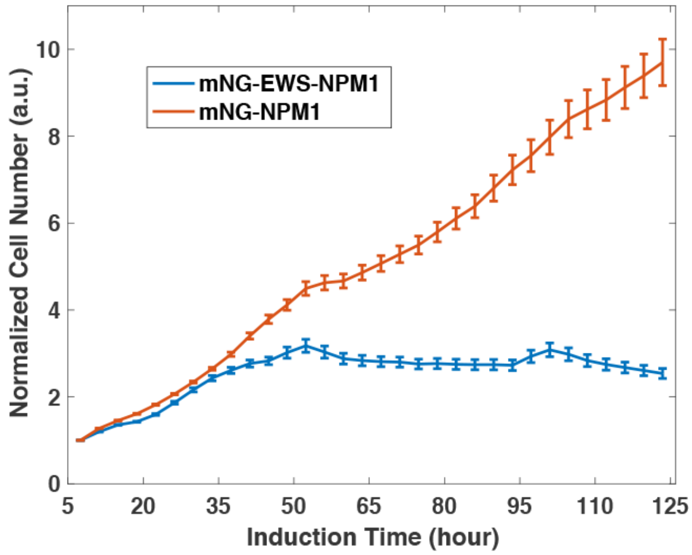


**Fig. S9. Cell number over time estimated using RTCA for knock-in A673 cells after induction of mNG-EWS-NPM1 (blue) or mNG-NPM1 (orange).**

Each cell number is normalized to the cell number 7 hours after induction. Error bars represent standard deviations.

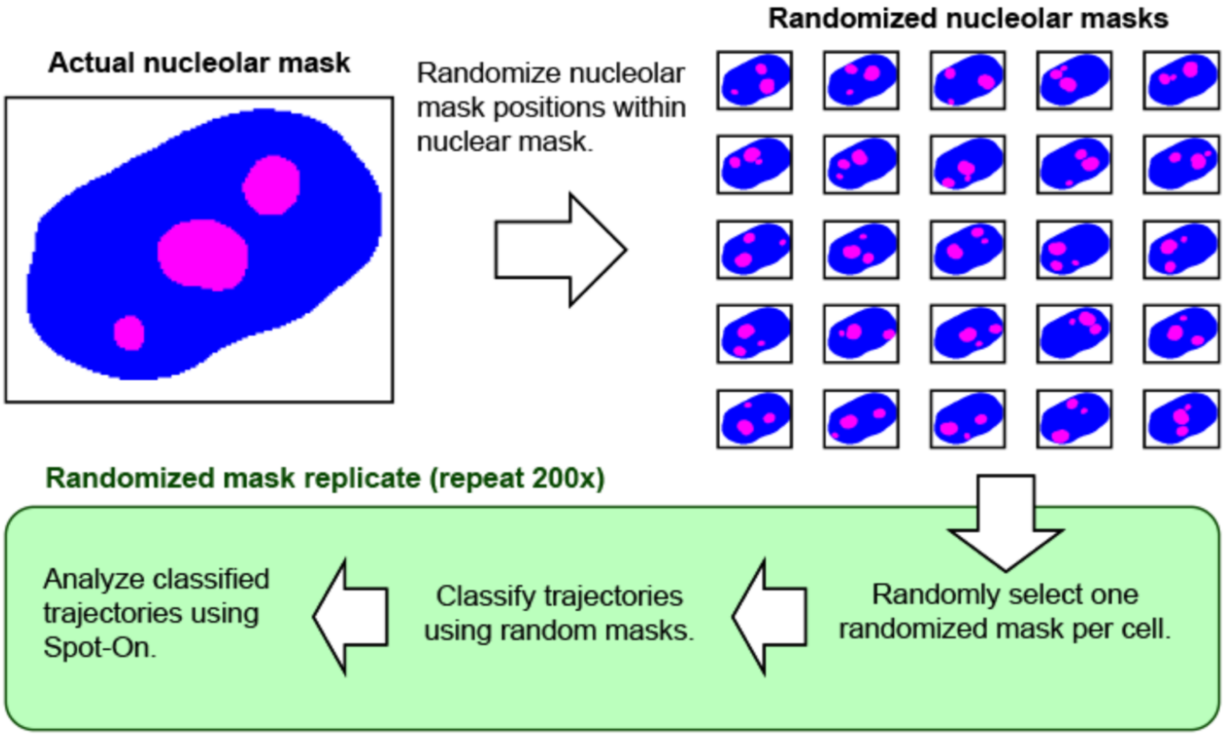


**Fig. S10. Procedure to analyze single-molecule trajectories of EWS/FLI1 using randomized nucleolar masks.**

Classifying trajectories using randomized masks provides a control for bias in diffusion analysis due to boundary effects of finite-sized ROIs.

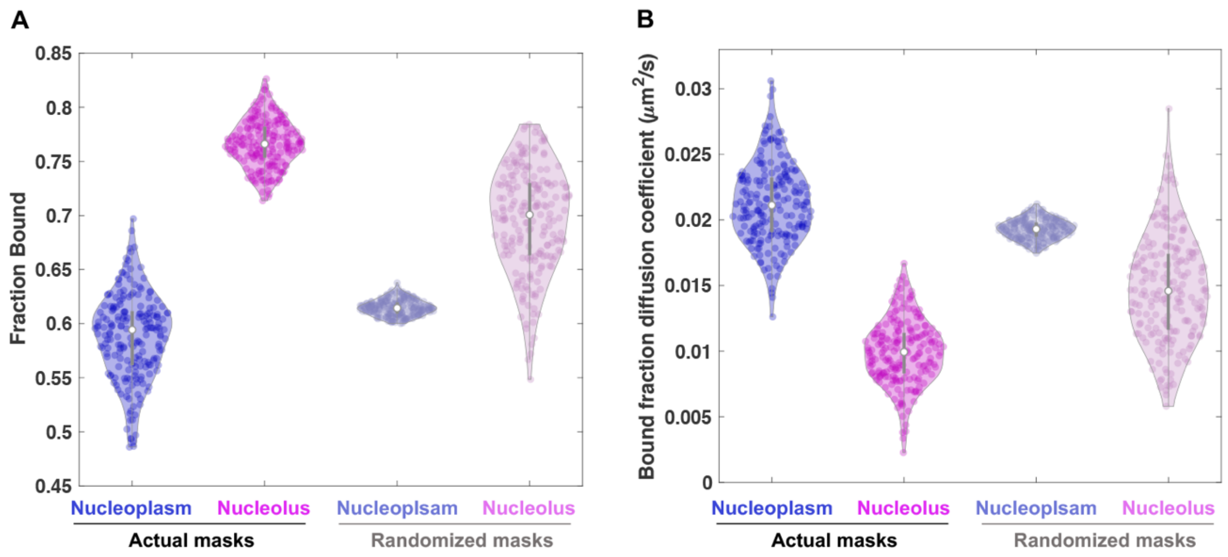


**Fig. S11. Fitted fraction of bound molecules (A) and diffusion coefficient of bound molecules (B) in the nucleoplasm (blue) and nucleolus (magenta).**

Left 2 columns: Fits for trajectories classified using actual nucleolar masks. Violin plots show the distribution of 200 bootstrap replicates resampled by cell with replacement. Right 2 columns: Fits for trajectories classified using 200 distinct sets of randomized nucleolar masks (see Supplemental Methods).

**Supplemental References:**


53. L. N. Randolph, X. Bao, C. Zhou, X. Lian, An all-in-one, Tet-On 3G inducible PiggyBac system for human pluripotent stem cells and derivatives. *Sci. Rep.* **7**, 1549-6 (2017).
54. I. Solovei, M. Cremer, 3D-FISH on cultured cells combined with immunostaining. *Methods Mol. Biol.* **659**, 117-126 (2010).
55. M. Tokunaga, N. Imamoto, K. Sakata-Sogawa, Highly inclined thin illumination enables clear single-molecule imaging in cells. *Nat. Methods* **5**, 159-161 (2008).
